# Decoding the physical principles of two-component biomolecular phase separation

**DOI:** 10.1101/2020.08.24.264655

**Authors:** Yaojun Zhang, Bin Xu, Benjamin G. Weiner, Yigal Meir, Ned S. Wingreen

**Affiliations:** Center for the Physics of Biological Function, Princeton University; Department of Physics, Princeton University; Department of Physics, Ben Gurion University of the Negev, Department of Physics, Princeton University and Department of Molecular Biology, Princeton University; Department of Molecular Biology, Princeton University and Lewis-Sigler Institute for Integrative Genomics, Princeton University

## Abstract

Cells possess a multiplicity of non-membrane bound compartments, which form via liquid-liquid phase separation. These condensates assemble and dissolve as needed to enable central cellular functions. One important class of condensates is those composed of two associating polymer species that form one-to-one specific bonds. What are the physical principles that underlie phase separation in such systems? To address this question, we employed coarse-grained molecular dynamics simulations to examine how the phase boundaries depend on polymer valence, stoichiometry, and binding strength. We discovered a striking phenomenon – for sufficiently strong binding, phase separation is suppressed at rational polymer stoichiometries, which we termed the magic-ratio effect. We further developed an analytical dimer-gel theory that confirmed the magic-ratio effect and disentangled the individual roles of polymer properties in shaping the phase diagram. Our work provides new insights into the factors controlling the phase diagrams of biomolecular condensates, with implications for natural and synthetic systems.

## I. INTRODUCTION

Eukaryotic cells are host to a multiplicity of non-membrane bound compartments. Recent studies have shown that these compartments form via liquid-liquid phase separation [1–3]. The phase-separated condensates enable many central cellular functions – from ribosome assembly, to RNA regulation and storage, to signaling and metabolism [2, 4]. Unlike conventional liquid-liquid phase separation, e.g. water-oil demixing, the underlying interactions that drive biomolecular phase separation typically involve strong one-to-one saturable interactions, often among multiple components. As a result, the phase diagrams of biomolecular condensates are complex and are sensitive to a variety of physical properties of the biomolecules, included number of binding sites, binding strengths, and additional non-specific interactions. Importantly, these physical parameters can be subject to biological regulation, and can thus directly impact the organization and function of the condensates. It is therefore crucial to understand how the physical properties of the components shape the phase diagram of biomolecular condensates.

Biomolecular condensates typically contain tens to hundreds of types of molecules. Yet, when characterized in detail, only a small number of components are responsible for condensate formation [4]. One class of such condensates are those formed by the association of two essential components. In the simplest case, each component consists of repeated domains that bind in a one-to-one fashion with the domains of the other component. Such two-component condensates have been observed in several natural and engineered contexts, including the algal pyrenoid, P-bodies, and promyelocytic leukemia (PML) nuclear bodies [5–7].

Previous simulations [5, 8] of average cluster size in such two-component systems revealed a striking phenomenon – for sufficiently strong binding, the formation of large clusters is suppressed when the valence of one species equals or is an integral multiple of the valence of the other species, favoring the formation of small stable oligomers instead of a condensate. The phenomenon reminiscent of the exact filling of atomic shells leading to the unreactive noble gases was termed the “magic-number” effect. However, as cluster size is subject both to finite-size effects and to ambiguity between a thermodynamic phase transition and a sol-gel percolation transition [9], it provides at best a qualitative measure of phase separation. Moreover, the previous studies focused on equal monomer stoichiometry, whereas biomolecular condensates cover a broad range of stoichiometries both *in vitro* [6, 7] and *in vivo* [10].

Here, we directly delineate the full phase diagram of such two-component systems. Using coarse-grained molecular dynamics simulations, we explore systematically how phase boundaries depend on valence, stoichiometry, and binding strength of two associating polymers. Our studies reveal an unanticipated effect – when the numbers of polymers of the two types have a rational stoichiometry (1:1, 1:2, etc.) and their valences are such that small oligomers can form with few unsatisfied bonds, then phase separation is strongly suppressed, which we call the “magic-ratio” effect. To understand the magic-ratio effects better, we develop a two-component sticker theory à la Semenov and Rubinstein [11, 12]. We model the system as dominated by polymer dimers in the dilute phase and by a condensate of independent stickers in the dense phase. The resulting analytical theory captures the magic-ratio effect discovered in simulations, and allows us to disentangle the individual roles of valence, stoichiometry, specific-bond strength, and non-specific attraction in determining the phase boundaries of two-component multivalent systems. Living cells regulate the valence and interactions of biomolecules through chemical modification, or on a slower timescale, tune the stoichiometry via synthesis/degradation or sequestration, and over evolutionary time, adapt the strength of specific and non-specific interactions through mutation of molecular sequences. Understanding the individual roles of these biologically tunable variables thus brings new insights into possible cellular strategies for regulating the formation and dissolution of biomolecular condensates.

## II. SIMULATION RESULTS

We perform coarse-grained molecular-dynamics simulations using LAMMPS [13] to determine the phase boundaries of two-component multivalent systems (Fig. 1). Briefly, we model the two polymer species as flexible linear chains of beads connected by harmonic springs (Fig. 1*A*). Each bead represents one associative domain of the polymer. To ensure associative domains of different polymer types bind in a one-to-one fashion, we impose a finite-ranged attractive interaction between beads of different types. This, however, could lead to more than one-to-one associations. Therefore, to avoid such unwanted associations, we impose strong repulsive interactions between beads of the same type over a large enough range to prevent other beads overlapping with a bound pair, thus preventing multiple-to-one binding (Figs. 1*B* and S1), see Methods and Supplemental Material for details.

**FIG. 1.**
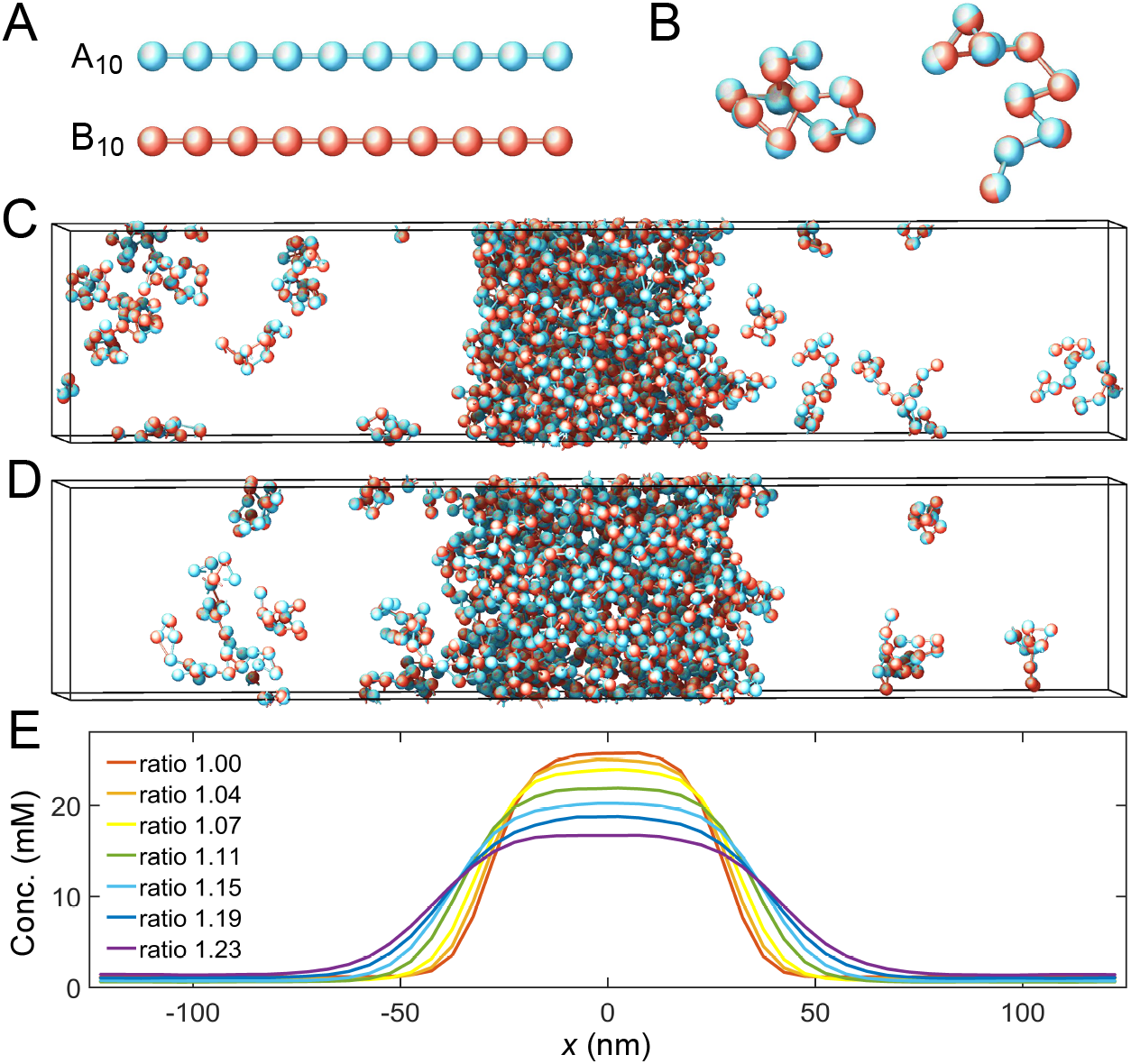
Coarse-grained molecular-dynamics simulations of two-component multivalent associative polymers. (*A*) The system consists of two types of polymers A (blue) and B (red) of varying lengths and concentrations. Depicted are A and B polymers of length 10, denoted as A_10_ and B_10_. Each polymer is modeled as a linear chain of spherical particles connected by harmonic bonds. Monomers of different types interact pairwisely through an attractive potential, while repulsion between monomers of the same type prevents them from overlapping and thus ensures one-to-one binding of monomers of different types (see Methods and Supplemental Material for details). (*B*) Snapshots of dimers formed by A_10_ and B_10_ with one-to-one bonds. (*C*) Snapshot of a simulation with 125 A_10_ and 125 B_10_ polymers. The system phase separates into a dense phase (middle region) and a dilute phase (two sides) in a 250nm×50nm×50nm simulation box with periodic boundary conditions. (*D*) Same as *C* but with 138 A_10_ and 112 B_10_ polymers, yielding an overall monomer concentration ratio (stoichiometry) 1.23. (*E*) Monomer concentration profiles of A_10_:B_10_ systems at various overall stoichiometries (total monomer concentration 6.64mM), each with the center of the dense phase aligned at *x* = 0 and averaged over time and over ten simulation repeats (see Methods and Supplemental Material). All simulations performed in LAMMPS [13].

To find the phase boundaries, we simulate hundreds of polymers of types A and B with, respectively, *m* and *n* monomers (an A_*m*_:B_*n*_ system) in a box with periodic boundary conditions (Figs. 1*C* and 1*D*). We initialize the system by constructing a dense slab of polymers in the middle of the box [14]. The system evolves and relaxes according to Langevin dynamics [15]. After the system has achieved equilibrium, two phases coexist: a dilute phase consisting of dimers and other small oligomers, and a dense phase of an interconnected polymer condensate. We measure the corresponding density profile (Fig. 1*E*) and calculate the dilute- and dense-phase concentrations by averaging the density profile over the regions (*x* ≤ −100nm or *x* ≥ 100nm) and (−10nm ≤ *x* ≤ 10nm), respectively. See Methods and Supplemental Material for simulation details.

### A. Equal Monomer Stoichiometry

It was shown previously that for equal monomer stoichiometry in the strong-binding regime, clustering is substantially suppressed when the number of binding sites on one polymer species is an integer multiple of the number of binding sites on the other, as this condition favors the assembly of small oligomers in which all binding sites are saturated [5, 8]. What does this magic-number effect imply for the actual phase diagram? To address this question, we fix the valence of polymer A at 14 and systematically vary the valence of polymer B from 5 to 16 while keeping the two monomer concentrations the same, i.e., at equal global monomer stoichiometry.

Figures 2*A* and 2*B* show simulation results for total monomer concentrations of the dilute and dense phases for A_14_:B_5_ to A_14_:B_16_ systems. In the strong binding regime, for magic-number cases, i.e. when the valence of B is 7 or 14, the dilute-phase concentration shows pronounced peaks (Fig. 2*A*, black curve). What is the origin of the peak at A_14_:B_14_? Intuitively, when the dilute phase of the two-component system is dominated by dimers (for systems A_14_:B_12_ to A_14_:B_16_, as supported by cluster size analysis in Fig. S2), each of these dimers has high translational entropy, whereas polymers in the dense condensate have low translational entropy. For A_14_:B_14_, all binding sites can pair up in a dimer just as well as in the condensate, so the energy per polymer is not necessarily lower in the condensate. Why then is the condensate still competitive with the dilute phase? In a dimer, the binding sites of A_14_ must match all the binding sites of B_14_, leading to a reduced overall conformational entropy. By comparison, the polymers in the condensate are more independent, binding to multiple members of the other species and enjoying a relatively higher overall conformational entropy. Because the translational entropy of each dimer decreases as their concentration goes up, the condensed phase eventually becomes more favorable and so the system phase separates with increasing concentration. Therefore, phase separation in A_14_:B_14_ is primarily driven by a competition between translational entropy and conformational entropy.

**FIG. 2.**
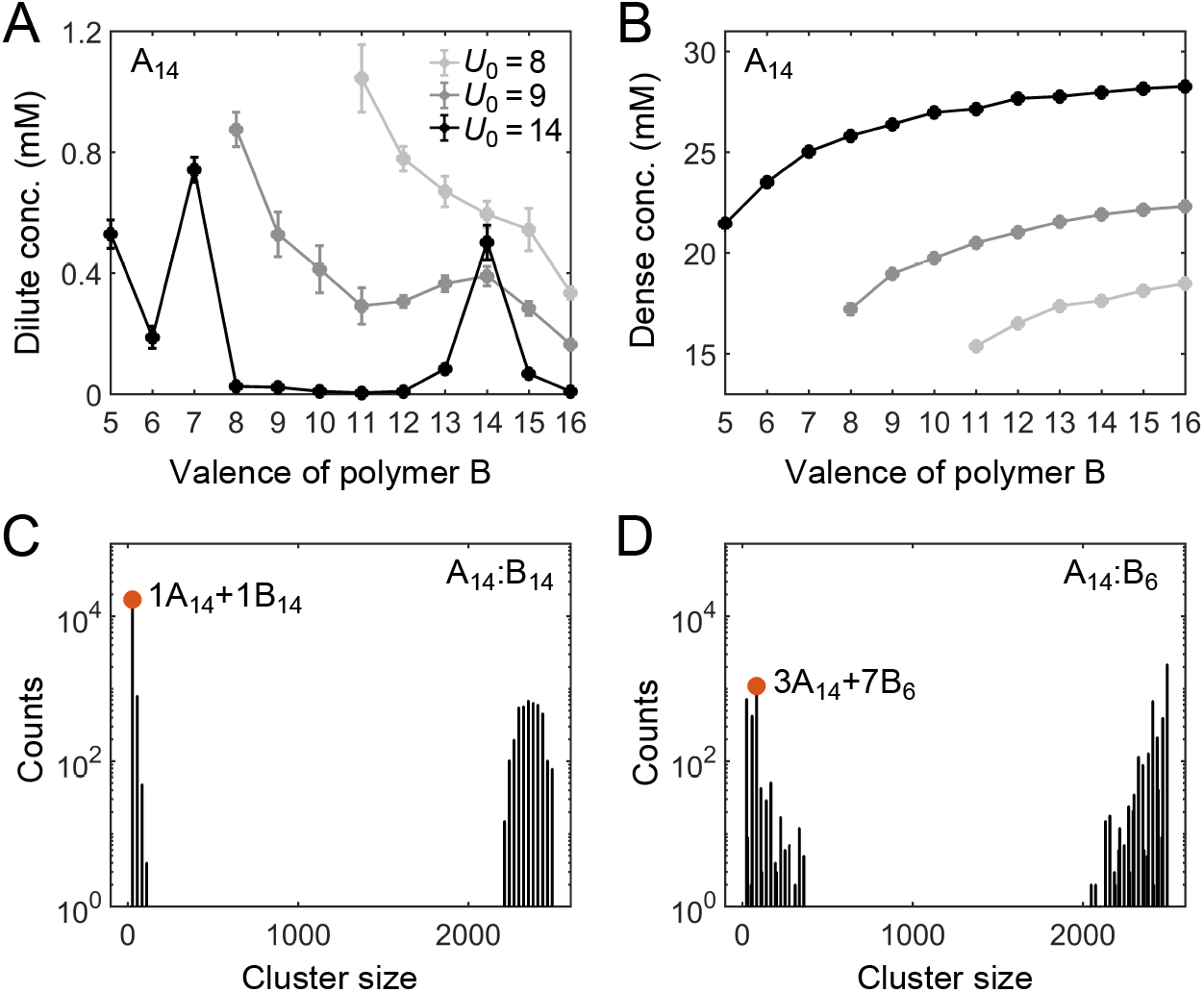
Simulations of associative polymers reveal a magic-number effect with respect to relative valence. Monomer concentrations in (*A*) dilute and (*B*) dense phases for simulated polymer systems at different binding strengths. *U*_0_ denotes the depth of the potential well, in units of *k*_B_*T* (see Methods and Supplemental Material for details). The valence of polymer A is 14, and the valence of polymer B ranges from 5 to 16. Global monomer stoichiometry is 1 and total monomer concentration is 6.64mM. Histograms of cluster size in (*C*) A_14_:B_14_ and (D) A_14_:B_6_ systems, for *U*_0_ = 14. “Counts” refer to number of clusters. Cluster size is measured in monomers. Red dots indicate the dominant oligomer in the dilute phase.

By contrast, for A_14_:B_13_ and A_14_:B_15_, one of the monomers in the dimer cannot be paired, and for A_14_:B_12_ and A_14_:B_16_, two monomers per dimer cannot be paired. Therefore, forming a condensate not only increases the conformational entropy but more importantly lowers the energy of these systems. This significantly tilts the balance in favor of condensation. As a result, the dilute-phase concentration is sharply peaked at A_14_:B_14_, falling off rapidly for increasingly unequal polymer lengths. We note that the dense-phase concentration shows no such feature (Fig. 2*B*), indicating that the peak at A_14_:B_14_ does not arise from differences in the internal structure of the dense phase.

The dilute phase of two-component systems is not always dominated by dimers (Fig. S2). For example, the dilute phase of the A_14_:B_7_ system is dominated by fully-bonded trimers with 1 A_14_ and 2 B_7_, the dilute phase of A_14_:B_8_ is dominated by trimers with 1 A_14_ and 2 B_8_, which has two unpaired monomers per trimer, and the dilute phase of A_14_:B_6_ is dominated by oligomers with 3 A_14_ and 7 B_6_, which although fully-bonded is not small (Fig. 2*D*). Consistent with the above logic, we find another peak in the dilute-phase concentration at A_14_:B_7_ (Fig. 2*A*). More generally, in contrast to the magic-number systems, the dilute phases in other cases are dominated by oligomers which are not capable of being fully bonded (high energy) and/or not small (low translational entropy) (Fig. S2). The dilute-phase concentration is therefore lower in these non-magic-number cases.

How do the phase boundaries depend on the strength of binding? Figure 2*A* shows that, for non-magic-number systems, the dilute-phase concentration decreases monotonically with increasing binding strength, whereas for magic-number systems the dependence can be non-monotonic. This difference is attributed to the distinct underlying driving forces for phase separation. For non-magic-number systems, as clustering allows a larger fraction of binding sites to be paired, the stronger the binding, the more the energy is lowered by condensate formation. Therefore, the dilute-phase concentration drops as binding strength increases (or as temperature decreases). Such energy-dependence is expected for conventional phase-separation models, such as Flory-Huggins [16, 17].

Interestingly, for the magic-number system A_14_:B_14_, the dilute-phase concentration first decreases with increasing binding strength in the weak binding regime, similar to non-magic-number systems. However, as the binding energy is increased further, most of binding sites pair up in both dilute and dense phases. Phase separation is then primarily driven by a competition between conformational and translational entropy. The pairing up of binding sites reduces the conformational entropy of both the dense and dilute phases. By contrast, the translational entropy of the dilute-phase components is almost unaffected. Consequently, the dilute phase becomes more competitive relative to the condensate, so the dilute phase boundary shifts to higher concentration.

By comparison, the dense-phase concentration increases monotonically with increasing binding strength for all systems (Fig. 2*B*). This follows because the stronger the binding, the more monomers are paired, which tightens the condensate structure. We note that at substantially higher binding energies than studied here, essentially all the binding sites are satisfied in both magic-number and non-magic-number systems, and the phase boundaries become independent of binding energy.

### B. Unequal Monomer Stoichiometry

How do the phase boundaries depend on overall monomer stoichiometry? Figures 3*A* and 3*B* show total monomer concentrations of the dilute and dense phases for magic-number systems A_8_:B_8_ to A_14_:B_14_ at different global monomer stoichiometries. For each system, the dilute-phase concentration peaks at equal monomer ratio, falls off initially as the ratio deviates from 1, and then curves back up. What is the origin of the peak at equal stoichiometry? Recall that, in the strong binding regime, phase separation of magic-number systems is primarily driven by a competition between translational entropy and conformational entropy. Now consider starting with a system at equal monomer concentration, and adding more of one polymer species to the system. At the beginning, the added polymers readily enter the dense phase, which relaxes the conformational constraint that every monomer in the condensate has to pair with a partner. This increase of the conformational entropy of the condensate makes it more competitive, so the dilute-phase concentration decreases. However, as the ratio between the two polymers is increased further, it becomes possible to form a spectrum of dilute-phase oligomers which typically contain one extra polymer of the majority type (Table S1). These new oligomers have more relaxed structures than fully-bonded dimers, which raises the conformational entropy of the dilute phase. Therefore, the dilute phase is favored over the condensate and its concentration curves back up.

**FIG. 3.**
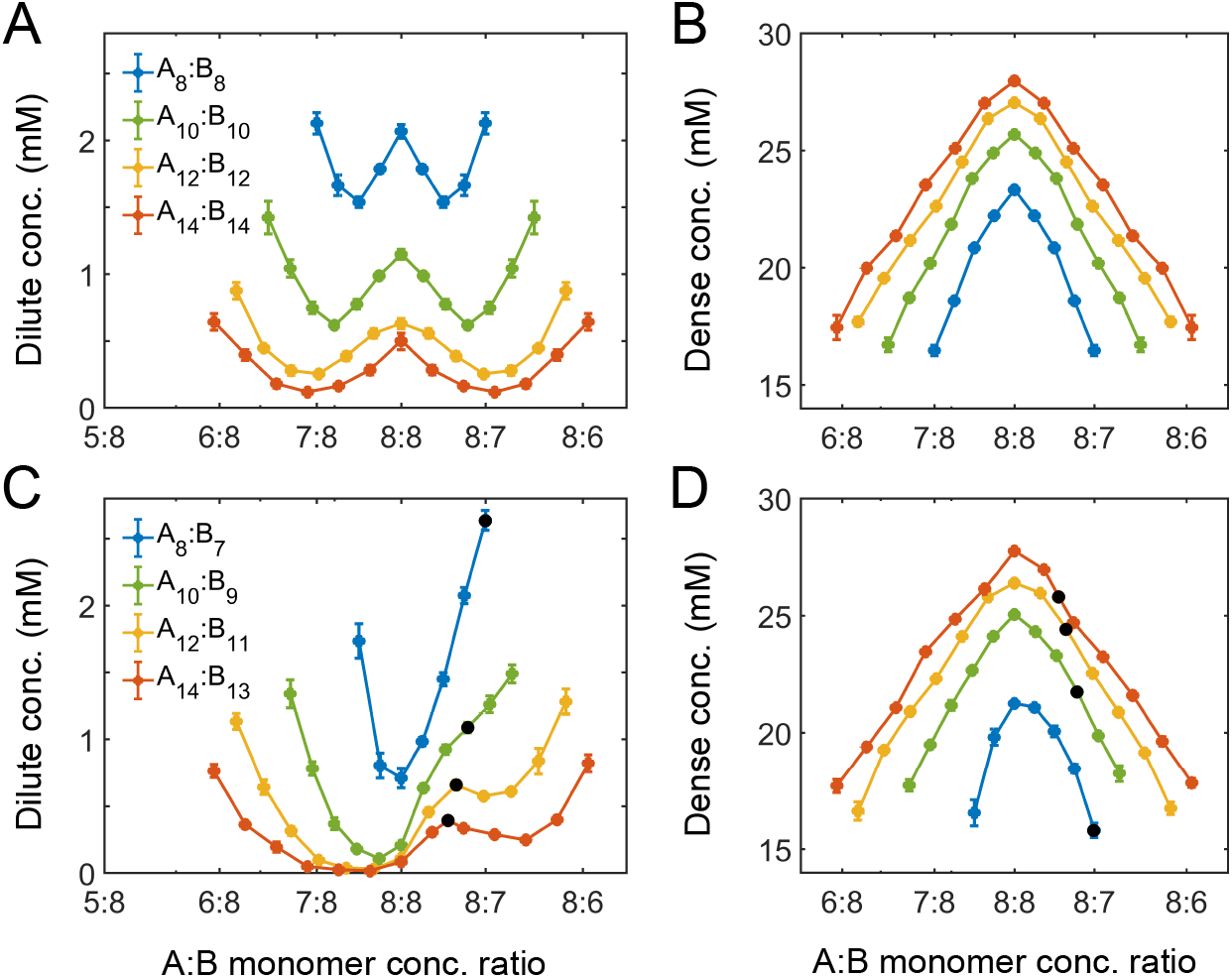
Simulations of associative polymers reveal a magic-ratio effect with respect to polymer stoichiometry. Monomer concentrations in (*A*) dilute and (*B*) dense phases for equal polymer length systems (i.e., A_*n*_:B_*n*_) at different global stoichiometries. Monomer concentrations in (*C*) dilute and (*D*) dense phases for systems where polymer B is 1 monomer shorter than polymer A (i.e., A_*n*_:B_*n*–1_) at different stoichiometries; black dots indicate cases where the number of polymers of each type is the same. Interaction strength *U*_0_ = 14 and total monomer concentration is 6.64mM.

Figure 3*A* also reveals that the dilute-phase concentration decreases with increasing polymer valence. This follows in part because translational entropy in the dilute phase is per dimer center of mass, whereas conformational entropy in both phases scales with the number of monomers. The entropic gain of joining the dense phase is therefore more on a per monomer basis for longer polymers, so the dilute-phase concentration decreases with increasing valence. As a less apparent yet important point, Fig. 3*A* also shows that increasing polymer valence enhances both the width and relative height of the peak in the dilute-phase concentration.

Figures 3*C* and 3*D* show total monomer concentrations of the dilute and dense phases for un-equal valence polymers A_8_:B_7_ to A_14_:B_13_ at different global monomer stoichiometries. The dilute phase boundary shows a symmetric minimum around equal stoichiometry for A_8_:B_7_, yet surprisingly, the phase boundary becomes asymmetric and then peaks at equal *polymer* stoichiometry with increasing polymer length (Fig. 3*C*). What is the origin of these peaks? Taking the A_14_:B_13_ system as an example, its dilute phase is dominated by dimers with an unpaired A monomer. This strongly disfavors the dilute phase in the strong binding regime at equal monomer stoichiometry. However, as the overall A:B monomer stoichiometry increases, the excess As cannot be paired anyway. In particular, at equal polymer stoichiometry (denoted as black dots in Fig. 3*C*), forming dimers is no longer energetically costly. The result is strong suppression of phase separation at equal polymer stoichiometry. We term this the ”magic-ratio” effect, as it occurs for rational ratios of associative polymers with strong one-to-one bonds. The effect is enhanced for longer polymers with few unsatisfied bonds, hence the clear peak in the dilute phase concentration for the A_14_:B_13_ system. Taking this case as an example, we can simply understand why the peak occurs at equal polymers stoichiometry. To the left of the A_14_:B_13_ peak at equal polymer stoichiometry, the dilute-phase concentration is low as dimers are energetically disfavored because more bonds can be satisfied in the condensate. By contrast, to the right of the peak, the dilute-phase concentration is low for a different reason – because the condensate is entropically favored, similar to the peak with respect to stoichiometry for magic-number systems. Eventually, the dilute-phase concentration curves back up due to formation of higher oligomers in the dilute phase, as discussed for magic-number systems.

We note that for all these systems the dense-phase concentration shows no such striking features. Rather, the concentration decreases monotonically as the global monomer stoichiometry departs from 1 and as the valence of polymers decreases (Figs. 3*B* and 3*D*).

Above, we considered the role of both relative valence and relative stoichiometry. By plotting phase boundaries as joint functions of valence and stoichiometry, we obtain a unified picture: Figures 4*A* and 4*B* show the dilute- and dense-phase concentrations for systems A_14_:B_12-16_ at global monomer stoichiometries 14:12-16. Notably, the dilute-phase concentration is peaked along the diagonal (Fig. 4*A*), i.e. at equal *polymer* stoichiometry, which we termed the “magic-ratio effect”. Intuitively, all cases along the diagonal favor 1:1 polymer dimers: the dimers enjoy high translation entropy and there is no energy penalty involved in their formation. Thus, a dilute phase of dimers is strongly favored at equal polymer stoichiometry.

**FIG. 4.**
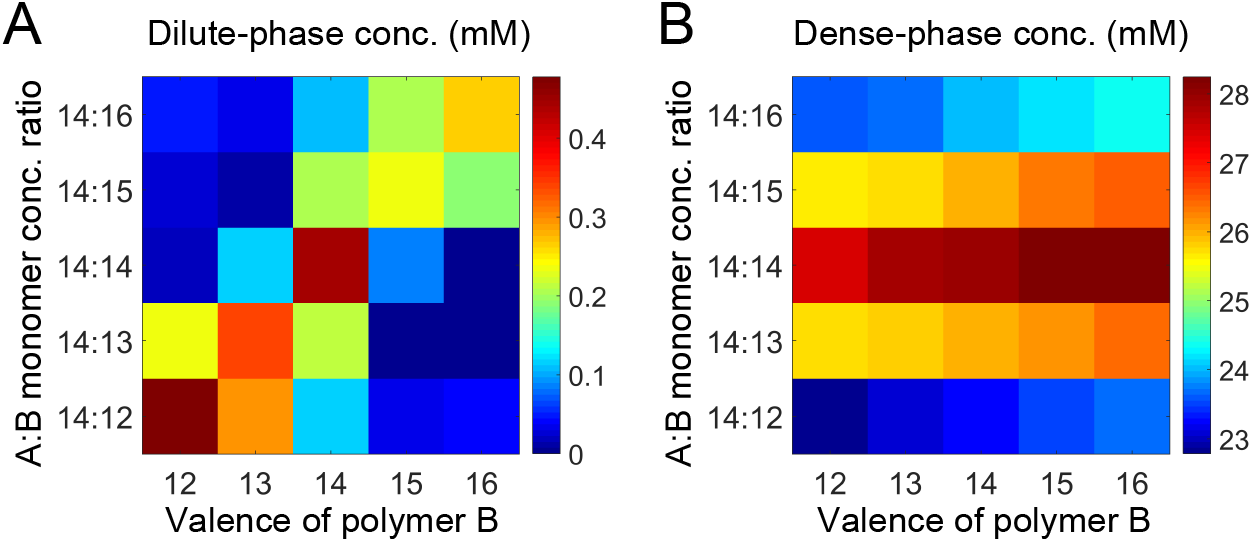
Simulations of associative polymers reveal a magic-ratio effect with respect to polymer stoichiometry. Monomer concentrations in (*A*) dilute and (*B*) dense phases for systems A_14_:B_12-16_ at global stoichiometries 14:12-16. Interaction strength *U*_0_ = 14 and total monomer concentration is 6.64mM.

## III. DIMER-GEL THEORY RESULTS

While our simulations have revealed that a magic-ratio effect influences the boundaries of phase separation for associating polymers, we desire a deeper understanding of the interplay of factors such as overall valence, stoichiometry, and interaction strength. To this end, we develop a mean field theory of two-component associative polymers à la Semenov and Rubinstein [11, 18].

Specifically, we consider a system of A and B polymers as in our simulations. Each polymer is a linear chain of *L*_1_ or *L*_2_ monomers of type A or type B, respectively. Without loss of generality, we take *L*_1_ ≥ *L*_2_. Monomers (stickers) of different types associate in a one-to-one fashion. Our simulations suggest that for polymers of similar valence close to equal polymer stoichiometry the dilute phase is dominated by dimers and the dense phase is a condensate. Therefore, we assume that polymers can associate either as dimers or, alternatively, as a condensate in which pairs of monomers bind independently. This assumption of independence is a mean field approximation, as it neglects correlations between monomers in the same chain, and thus only applies when the polymers strongly overlap, i.e. at densities above the semidilute regime [19].

The partition function of such a system can be divided into three parts: *Z* = *Z*_ni_*Z*_s_*Z*_ns_, where *Z*_ni_, the partition function of a solution of non-interacting polymers, captures the translational and conformational entropy of the two polymer species, *Z*_s_ captures specific interactions between associating monomers, and *Z*_ns_ captures all non-specific interactions.

The corresponding free-energy density for the mixed non-interacting polymers is [11]

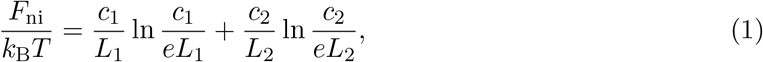

where *c*_1_ and *c*_2_ are the concentrations of A and B polymers measured in terms of monomers. Note that the terms for the conformational entropy of non-interacting polymers are omitted in Eq. (1), as they are linear in *c*_1_ and *c*_2_ and thus do not influence the phase boundaries.

To include specific interactions, we first consider the partition function *Z*_s_(*N*_d1_, *N*_d2_, *N*_b_) for states with exactly *N*_d1_ and *N*_d2_ total numbers of monomers of A and B types in dimers (i.e. number of dimers equals *N*_d1_/*L*_1_ = *N*_d2_/*L*_2_) and *N*_b_ additional monomer pairs,

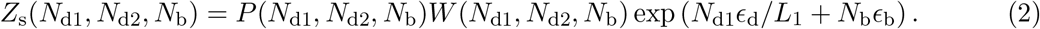

In Eq. (2), *P* is the number of different ways that polymers and monomers can be chosen to pair up to form dimers and independent bonds,

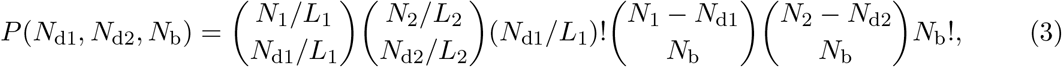

where *N*_1_ and *N*_2_ are the total numbers of monomers of A and B types. (Note that in Eq. (3) if *L*_1_ > *L*_2_, the excess monomers of type A in dimers do not form additional bonds.) In Eq. (2), *W* is the probability that all chosen polymers and monomers are, respectively, close enough to their specified partners in the non-interacting state to form dimers and independent bonds,

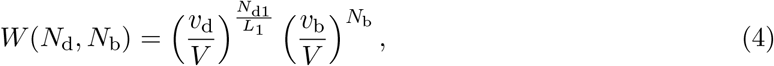

where *v*_d_ and *v*_b_ are effective interaction volumes and *V* is the system volume. The last term in Eq. (2) is the Boltzmann factor for specific interactions, where *ϵ*_d_ and *ϵ*_b_ are the effective binding energies of dimers and monomer pairs, in units of *k*_B_*T*.

The part of the free-energy density due to specific interactions is

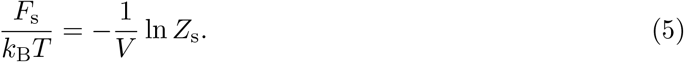

Using Stirling’s approximation ln *N*! = *N* ln *N* – *N*, we obtain

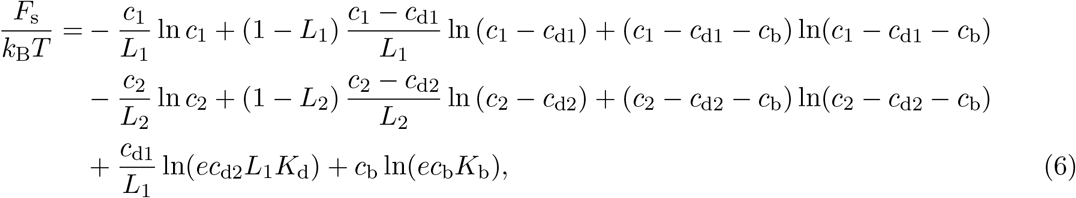

where *K*_d_ ≡ *e*^−*ϵ*_d_^/*v*_d_ and *K*_b_ ≡ *e*^−*ϵ*_b_^/*v*_b_ are, respectively, the dissociation constants of a dimer and of a pair of monomers. *c*_d1_ and *c*_d2_ are the concentrations of monomers of A and B types in dimers (so *c*_d1_/*L*_1_ = *c*_d2_/*L*_2_), and *c*_b_ is the concentration of independent bonds.

In the thermodynamic limit, *F*_s_ will be minimized with respect to *c*_d1_, *c*_d2_ and *c*_b_, which implies

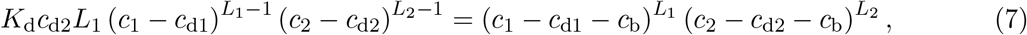

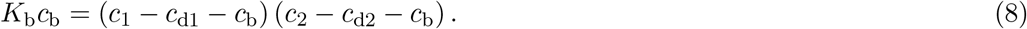

Note that if *c*_b_ in Eq. (7) and *c*_d1_ and *c*_d2_ in Eq. (8) are set to zero, these equations reduce to

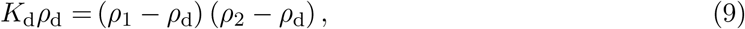

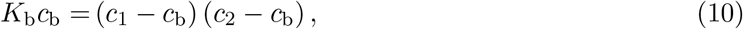

where *ρ*_1_, *ρ*_2_, and *ρ*_d_ are the total concentrations of A and B polymers and dimers (measured in polymeric units), i.e., *ρ*_1_ = *c*_1_/*L*_1_, *ρ*_2_ = *c*_2_/*L*_2_, and *ρ*_d_ = *c*_d1_/*L*_1_ = *c*_d2_/*L*_2_. Eqs. (9) and (10) are consistent with the definitions of the dissociation constants of a dimer and of an independent bond, respectively.

The free-energy density due to non-specific interactions can in general be written as a power expansion in the concentrations [11, 19],

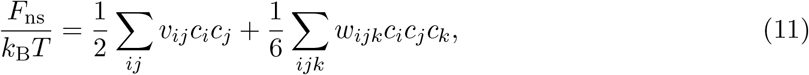

where the sum is over all the species in the system, including free polymers/monomers, dimers and independent bonds, and *v_ij_* and *w_ijk_* are two- and three-body interaction parameters. For our simulation system, we derive a specific form of *F*_ns_ by taking into account that (1) we are interested in the strong-binding regime where the magic-ratio effect is observed, (2) there is no non-specific interaction between free polymers of different types in our simulation, and (3) non-specific interactions are only important at high concentrations. The result is

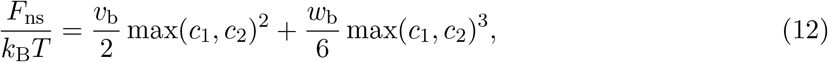

where *v_b_* and *w_b_* are the two- and three-body interaction parameters for a solution of independent bonds. See Appendix **??** for details of the derivation.

Finally, substituting the conditions Eqs. (7) and (8) into Eq. (6), we obtain the total free-energy density *F* = *F*_ni_ + *F*_s_ + *F*_ns_,

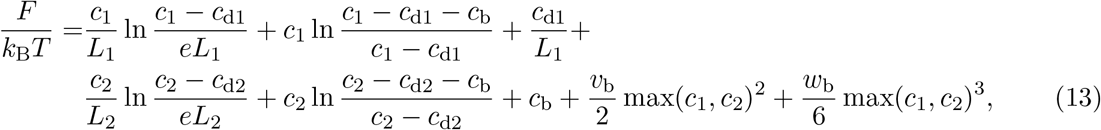

where *c*_d1_, *c*_d2_, and *c*_b_ are the solutions of Eqs. (7) and (8). Equations (7), (8), and (13) form a complete set which predicts the free-energy density of the two-component associative polymer system at given total monomer concentrations, *c*_1_ and *c*_2_, of the two species.

Intuitively, in the strong-binding regime, i.e. when *c*_1_, *c*_2_ ≫ *K*_d_, *K*_b_, polymers either associate as dimers or as independent bonds depending on their relative free energies. In the limit that dimers are preferred (*ρ*_d_ = min(*ρ*_1_, *ρ*_2_) and *c*_b_ = 0), the contribution from specific interactions is

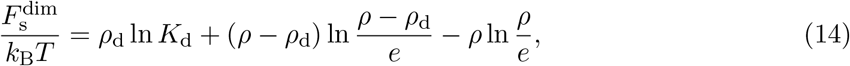

where *ρ* = max(*ρ*_1_, *ρ*_2_) is the concentration of the majority species in polymeric units. The terms on the right of Eq. (14) reflect, respectively, the free-energy density due to dimer formation, translational entropy of leftover polymers, and loss of translational entropy of the majority species (in effect, the formation of each dimer removes the translation entropy of one free polymer). In the opposite limit that independent bonds are preferred (*c*_b_ = min(*c*_1_, *c*_2_) and *ρ*_d_ = 0),

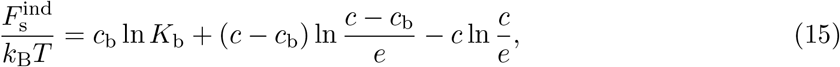

where *c* = max(*c*_1_, *c*_2_), and the terms are analogous to those in Eq. (14). Numerical studies show that the full *F*_s_(*c*_1_, *c*_2_) in Eq. (6) is always well approximated by the lower of the two limiting values of *F*_s_ (Eqs. (14) and (15)).

In which regions of concentration space are dimers versus independent bonds preferred? For a magic-number system composed of two polymer species of valence *L* at equal monomer stoichiometry, 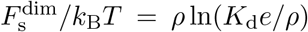 and 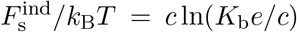. Comparing the two expressions, dimers are favored at low concentrations, whereas a gel connected by independent bonds is favored at high concentrations. The transition occurs when 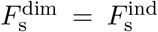, i.e. at concentration 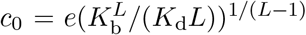. Away from equal stoichiometry, the transition occurs at a lower concentration *c_s_* = *c*_0_(*s* – 1)^*s*–1^*s*^−*s*^, where *s* = max(*c*_1_, *c*_2_)/ min(*c*_1_, *c*_2_) > 1 (see Supplemental Material for details). As *c_s_* decreases rapidly with increasing *s* (Fig. 5*A* inset, white curve), the preference for dimers over a gel exhibits a sharp peak around equal stoichiometry.

**FIG. 5.**
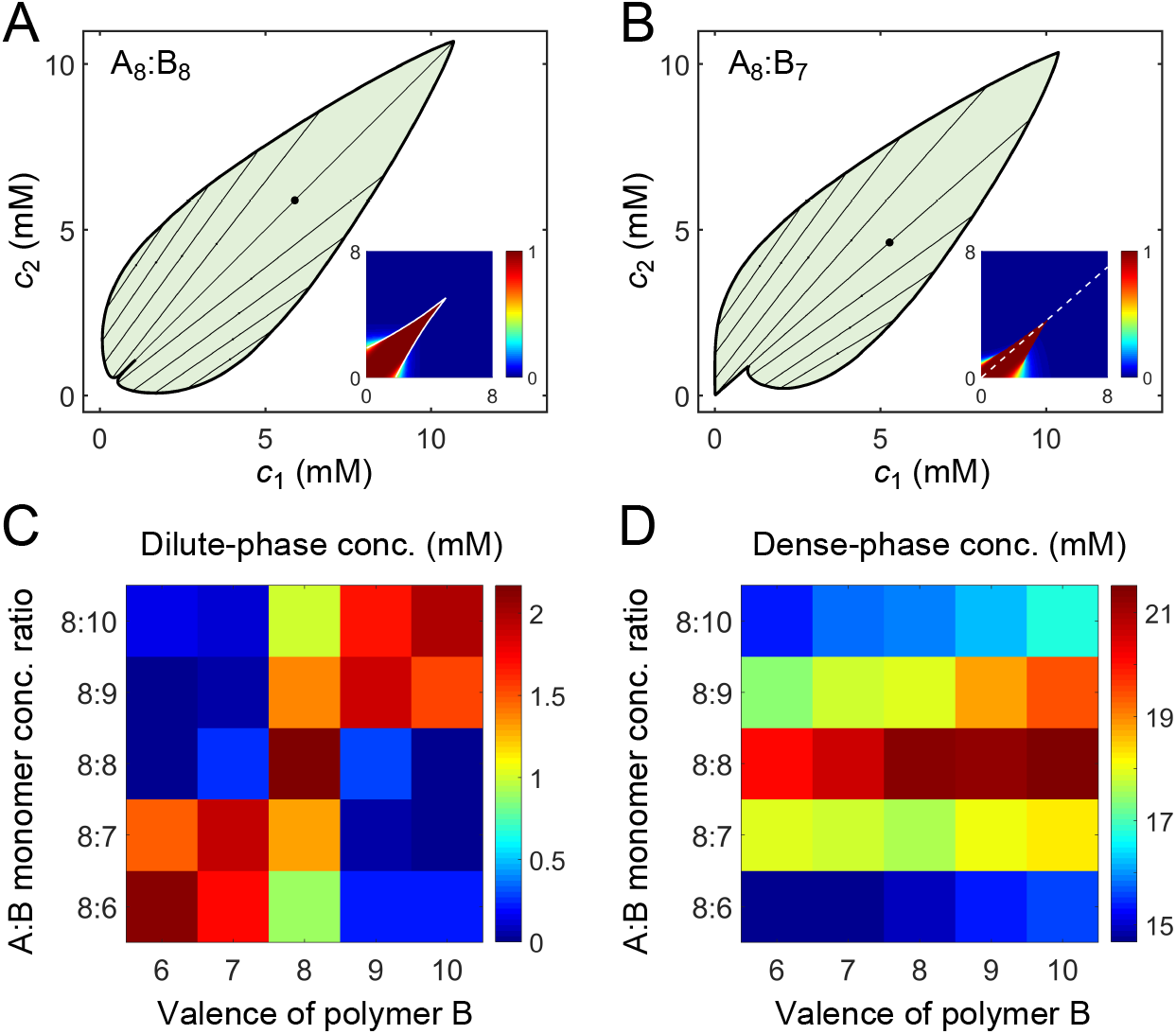
A dimer-gel theory predicts the magic-ratio effect. Phase diagrams of (*A*) A_8_:B_8_ and (*B*) A_8_:B_7_ systems: one-phase region white, two-phase region green. The dilute- and dense-phase concentrations are connected by representative tie lines. The tie line along the direction of equal polymer stoichiometry is denoted with a black dot. Insets: fraction of monomers in dimers for (*A*) A_8_:B_8_ and (*B*) A_8_:B_7_ systems. White curve in *A* inset is the transition boundary between dimer- and independent bonds-dominated regions predicted by *c_s_*. Dashed white line in (*B*) inset denotes equal polymer stoichiometry. Monomer concentrations in (*C*) dilute and (*D*) dense phases for systems A_8_:B_6-10_ at global monomer stoichiometries 8:6-10. The total monomer concentration is the same as in simulations, 6.64 mM. For details see Supplemental Material. Parameters: *v*_b_ = 9 × 10^−2^mM^−1^, *w*_b_ = 7 × 10^−3^mM^−2^, *K*_b_ = 3.8 × 10^−3^mM, and *K*_d_ values in Table S3.

To give a concrete example of the above analysis, we extract the values of *K*_d_ for dimers from simulations, choose a value of *K_b_* for independent bonds close to the dissociation constant of a pair of monomers (see Supplemental Material for details), and numerically solve Eqs. (7) and (8) for *c*_d1_, *c*_d2_, and *c*_b_ to find the fraction of monomers in dimers and independent bonds for all concentrations (*c*_1_, *c*_2_). We find that indeed for polymers of equal valence, dimers are favored at low concentrations and independent bonds at high concentrations. The dimer dominated region extends sharply to higher concentrations in a narrow zone around the diagonal, as quantitatively captured by *c_s_* (Figs. 5*A* inset and S3*A*). For polymers of similar but unequal valence, the dimer dominated region extends to higher concentrations along the direction of equal *polymer* stoichiometry (Figs. 5*B* inset and S3*B*).

Finally, to extract the phase boundaries, we substitute the values of *c*_d1_, *c*_d2_, and *c*_b_ into Eq. (13) to first obtain the free energy as a function of *c*_1_ and *c*_2_. The free-energy landscape has two basins, one at small concentrations corresponding to the dilute dimer-dominated phase, and one at high concentrations corresponding to the dense independent-bond-dominated gel-phase (Fig. S4). We locate the phase boundaries by applying convex-hull analysis to this free-energy landscape (see Supplemental Material).

Does the dimer-gel theory capture the magic-ratio effect revealed by our simulations? Figures 5*A* and 5*B* show the phase diagrams of A_8_:B_8_ and A_8_:B_7_ systems. In both cases, the phase boundaries on the dilute side extend sharply into the two-phase region along the direction of equal polymer stoichiometry (tie lines along this direction are denoted by black dots). Figures 5*C* and 5*D* show the dilute- and dense-phase concentrations for systems A_8_:B_6-10_ at global monomer stoichiometries 8:6-10. Notably, the dilute-phase concentrations are substantially shifted up around the diagonal, verifying the magic-ratio effect observed in simulations (Fig. 4*A*).

One of the major assumptions of the dimer-gel theory is the mean-field approximation, i.e. monomers of different types can associate independently in the dense phase. This approximation captures a key feature of the dense phase, namely that a single polymer binds to multiple partners. Nevertheless, because monomers belonging to the same polymer are tethered together, correlations in binding exist (Fig. S7). Therefore, what should be considered to be “independent” is not individual monomers but rather segments of the binding correlation length (~1.8 monomers). The dense phase of a valence 14 system is thus more accurately described by the theory at valence 14/1.8 ≈ 8. We therefore present results for valence 8 systems in Fig. 5. (The theoretical phase diagrams and the dilute- and dense-phase concentrations for valence 14 systems also verify the magic-ratio effect (Fig. S6).)

The dimer-gel theory has only a handful of parameters: the valences *L*_1_ and *L*_2_ of polymers A and B, the dissociation constants *K*_d_ and *K*_b_ of dimers and independent bonds, and the non-specific interaction parameters *v*_b_ and *w*_b_. How are the phase boundaries and the magic-ratio effects determined collectively by these parameters? If valence is increased while keeping all other parameters fixed in the theory, for equal valence polymers we find that the dilute-phase concentration decreases, while the dense-phase concentration increases, and the peak with respect to stoichiometry is enhanced in terms of the dilute-phase peak-to-valley ratio (Figs. S8*A* and S8*B*). If valence is increased for unequal valence polymers, we observe that the shape of the dilute phase boundary transitions from a shoulder to a peak (Figs. S8*C* and S8*D*). All these features are consistent with the simulation results in Fig. 3.

For the theory to agree quantitatively with the phase boundaries from simulations, we find that smaller values of non-specific interaction parameters are necessary for higher valence systems (Figs. S8*C* and S8*D*). Intuitively, this follows because higher valence polymers have more backbone bonds, which bring bound monomer pairs closer together in the dense phase – effectively reducing the non-specific repulsion between them. Finally, the dimer-gel theory also predicts that the magic-ratio effect disappears in the weak-binding regime (Fig. S9), consistent with our simulation results (Fig. 2).

## IV. DISCUSSION

Intracellular phase separation is driven by multivalent interactions between macromolecules. These interactions are separated into two classes [4, 20]: (1) specific interactions, such as binding between protein domains, are relatively strong and involve specific partners, and (2) non-specific interactions, such as electrostatic and hydrophobic interactions, which are much weaker, more generic, and non-saturable. Multivalent systems with specific interactions allow for “orthogonal” condensates to form: the specific interactions holding together one class of droplets will typically not interfere with those holding together another class. Motivated by the key role of specific interactions in intracellular phase separation, we focused on exploring the effects of specific interactions on the phase boundaries of two-component associative polymers. Specifically, we combined coarse-grained molecular dynamics simulations and analytical theory to examine the individual roles of valence, stoichiometry, and binding strength on the phase boundaries. In particular, we identified a magic-ratio effect: for sufficiently strong binding, phase separation is strongly suppressed at equal *polymer* stoichiometry.

The magic-ratio effect occurs exclusively in the strong-binding regime. Are specific protein-protein, protein-RNA, and RNA-RNA interactions strong enough to lead to the magic-ratio effect? The onset of the effect in our simulations occurs around *U*_0_ = 9*k*_B_*T* (Fig. 2*A*), which corresponds to a monomer-monomer dissociation constant *K*_d_ = 0.4mM. This value is consistent with the onset *K*_d_ of 1-2.5mM estimated from 3D lattice simulations with one polymer and one rigid component [8]. For comparison, the measured *K*_d_ for a SUMO protein domain with a SIM peptide is 0.01mM [7] and for an SH3 domain and a PRM peptide is 0.35mM [6]. Thus for systems as strongly interacting as SUMO-SIM or SH3-PRM, the magic-ratio effect in principle allows for novel mechanisms of regulation. For example, chemical modification of the effective valence of one component to change into or out of a magic-ratio condition could shift the phase boundary as a possible means of condensate regulation [5]. Magic-ratio effects could also manifest in other experimental systems, such as non-biological polymers, DNA origami [21], or patchy colloid systems [22]. As an inverse problem, the magic-ratio effect could be exploited to determine the relative valence of associating biomolecules by measuring their phase diagram.

While the magic-ratio effect is robust with respect to the strength of non-specific interactions, these interactions do strongly influence phase boundaries. For example, the dilute-phase concentrations in our simulations are ~1mM, while the reported values for biological systems are typically tens of *μ*M or less. The discrepancy is likely due to different strengths of non-specific attraction. Indeed, increasing the non-specific attraction in simulations from 0.06*k*_B_*T* to 0.12*k*_B_*T* leads to a 50% reduction in the dilute-phase concentration (Fig. S10). Future work will explore the interplay between specific and non-specific interactions, and their roles in determining the physical properties of droplets, such as surface tension, viscosity, etc.

The simulations and theory presented here are aimed at providing conceptual understanding. Quantitative descriptions of related real systems will likely require additional features, such as details of molecular shape and flexibility, linker lengths, as well as range and type of interactions. Nevertheless, we have discovered a universal magic-ratio effect, and provided evidence that it shapes the phase boundaries of multivalent associative polymers. We expect these general findings to be independent of most system details. We hope that our work will stimulate exploration of magic-ratio effects in both natural and synthetic multivalent, multicomponent systems.

## V. METHODS

### A. Modeling two-component multivalent associative polymers

We perform coarse-grained molecular-dynamics simulations using LAMMPS [13] to simulate two-component multivalent associative polymers. Individual polymers are modeled as linear chains of spherical particles connected by harmonic bonds (Fig. 1*A*). Harmonic bonds are modeled using a harmonic potential:

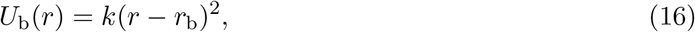

where *r*_b_ = 4.5nm is the mean bond length, 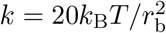 is the bond stiffness, *k*_B_ is the Boltzmann constant, and *T* = 300K is room temperature.

Monomers of the same type interact through a softened, truncated Lenard-Jones potential:

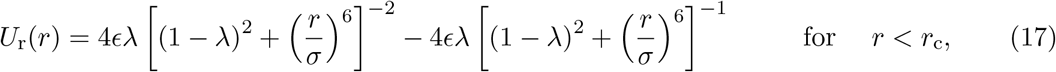

where *ϵ* = 0.15*k*_B_*T*, λ = 0.68, *σ* = 3.5nm, and *r*_c_ = 5nm. These parameters effectively lead to a monomer of diameter *d* ≃ 3nm and a weak attractive tail of depth 0.06*k*_B_*T*. The weak attractive tail is employed solely to promote a more compact condensate.

Monomers of different types interact through an attractive potential:

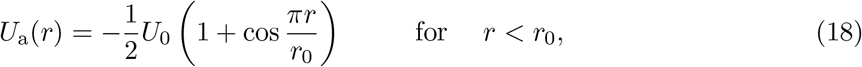

where *U*_0_ = 14*k*_B_*T* is used in all simulations, except as indicated in Fig. 2. The attraction cut-off distance is *r*_0_ = 2nm. Note that due to the strong repulsion between monomers of the same type, simultaneous binding of two monomers of one type to a monomer of the other type is energetically highly disfavored. This ensures one-to-one binding of monomers of different types (Figs. 1*B* and S1*D*).

### B. Determining the phase boundaries

Each system consists of *n*_1_ and *n*_2_ polymers of types A and B, respectively. The number of polymers are determined by their valences/lengths (*L*_1_ and *L*_2_) and global A:B monomer stoichiometry (*s*) through

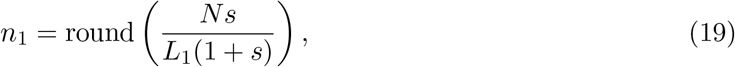

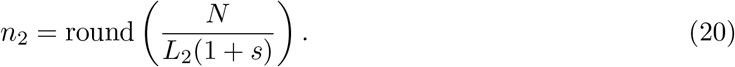

The round function is used as we can only simulate an integer number of polymers. For all simulations except those in Fig. 4, we use *N* = 2500 and a box of size 250nm×50nm×50nm. Simulations in Fig. 4 are performed with *N* = 5000 and a box of size 315nm×63nm×63nm.

Simulations are equilibrated using a Langevin thermostat in the *NVT* ensemble at *T* = 300K with periodic boundary conditions, i.e. the system evolves according to Langevin dynamics [15]

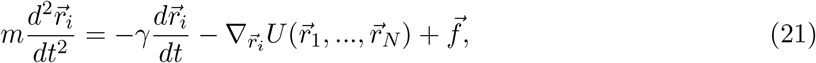

where 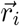 is the coordinate of particle *i, m* is its mass, *γ* is the friction coefficient, 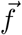 is random thermal noise, and the energy 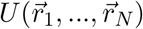 contains all interactions between particles, including harmonic bonds, non-specific, and specific interactions (Eqs. (16)–(18)).

To promote phase equilibrium and ensure that only a single dense condensate is formed, we first initialize the simulation by confining polymers in the region −50nm < *x* < 50nm. The attractive interaction between A and B monomers (Eq. (18)) is gradually switched on from *U*_0_ = 0 to 14 over 10^8^ time steps. This annealing procedure leads to the formation of a dense phase close to its equilibrated concentration. The confinement is then removed, and the system is equilibrated to allow the formation of a dilute phase and relaxation of the dense phase. We record the positions of all particles for 400 recordings after the system has achieved equilibrium. For each choice of valence and stoichiometry, we perform 10 simulation replicates with different random seeds.

To find the dilute- and dense-phase concentrations, we center the simulation box to the center of mass of the dense phase for each recording. We then compute the monomer concentration histogram along the *x* axis with a bin size 1/50 of box length. The resulting concentration profile has high values in the middle corresponding to the dense-phase concentration, and low values on the two sides corresponding to the dilute-phase concentration (Fig. 1*E*). The dilute- and dense-phase concentrations are calculated by averaging the concentration profile over the regions (*x* ≤ −100nm or *x* ≥ 100nm) and (−10nm ≤ *x* ≤ 10nm), respectively. Simulations with different total numbers of particles show consistent dilute- and dense-phase concentrations (Table S2), suggesting the effect of finite size is minor.

## VI. ACKNOWLEDGMENTS

We thank Guanhua He, Martin Jonikas, and Daniel Lee for insightful suggestions and comments. This work was supported in part by the National Science Foundation, through the Center for the Physics of Biological Function (PHY-1734030), and NSF grant 1521553.

## Notes

### Competing Interest Statement

The authors have declared no competing interest.

