## Supplemental Material for "Decoding the physical principles of two-component biomolecular phase separation"

### I. COARSE-GRAINED MOLECULAR-DYNAMICS SIMULATIONS

#### A. Modeling two-component multivalent associative polymers

We perform coarse-grained molecular-dynamics simulations using LAMMPS [1] to simulate two-component multivalent associative polymers. Individual polymers are modeled as linear chains of spherical particles connected by harmonic bonds (Fig. S1(a), type A polymer in blue and type B polymer in yellow). Harmonic bonds are modeled using a harmonic potential (Fig. S1(c), *left*)

$$U_{\rm b}(r) = k(r - r_{\rm b})^2, \tag{S1}$$

where $r_{\rm b} = 4.5$nm is the mean bond length, $k = 20k_{\rm B}T/r_{\rm b}^2$ is the bond stiffness, $k_{\rm B}$ is the Boltzmann constant, and $T = 300$K is room temperature.

Monomers of the same type interact through a softened, truncated Lenard-Jones potential (Fig. S1(c), *middle*) [8]

$$U_{\rm r}(r) = 4\epsilon\lambda \left\{ \left[ (1-\lambda)^2 + \left(\frac{r}{\sigma}\right)^6 \right]^{-2} - \left[ (1-\lambda)^2 + \left(\frac{r}{\sigma}\right)^6 \right]^{-1} \right\}, r < r_{\rm c}, \tag{S2}$$

where $\epsilon = 0.15k_{\rm B}T$, $\lambda = 0.68$, $\sigma = 3.5$nm, and $r_{\rm c} = 5$nm. These parameters effectively lead to a monomer of diameter $d \simeq 3$nm and a weak attractive tail of depth $0.06k_{\rm B}T$. The weak attractive tail is employed solely to promote a more compact dense condensate.

Monomers of different types interact through an attractive potential (Fig. S1(c), *right*)

$$U_{\rm a}(r) = -\frac{1}{2}U_0 \left(1 + \cos\frac{\pi r}{r_0}\right), r < r_0 \tag{S3}$$

where $U_0 = 14k_{\rm B}T$ is used in all simulations, except as indicated in Fig. 2. The attraction cut-off distance is $r_0 = 2$nm. Note that due to the strong repulsion between monomers of the same type, simultaneous binding of two monomers of one type to a monomer of the other type is energetically highly disfavored (Fig. S1(d)). This ensures one-to-one binding of monomers of different types (Fig. S1(b)).

#### B. Phase equilibration and data recording

Each system consists of $n_1$ and $n_2$ polymers of types A and B, respectively. The number of polymers are determined by their valences/lengths ($L_1$ and $L_2$) and global A:B monomer stoichiometry ($s$) through

$$n_1 = \text{round}\left(\frac{Ns}{L_1(1+s)}\right), \tag{S4}$$

$$n_2 = \text{round}\left(\frac{N}{L_2(1+s)}\right). \tag{S5}$$

The round function is used as we can only simulate an integer number of polymers. For all simulations except those in Fig. 4, we use $N = 2500$, so the total number of monomers is around 2500.

Simulations are equilibrated using a Langevin thermostat in the $NVT$ ensemble at $T = 300$K in a box of size 250nm×50nm×50nm with periodic boundary conditions, i.e. the system evolves according to Langevin dynamics [2]

$$m\frac{d^2\vec{r}_i}{dt^2} = -\gamma\frac{d\vec{r}_i}{dt} - \nabla_{\vec{r}_i}U(\vec{r}_1, ..., \vec{r}_N) + \vec{f}. \tag{S6}$$

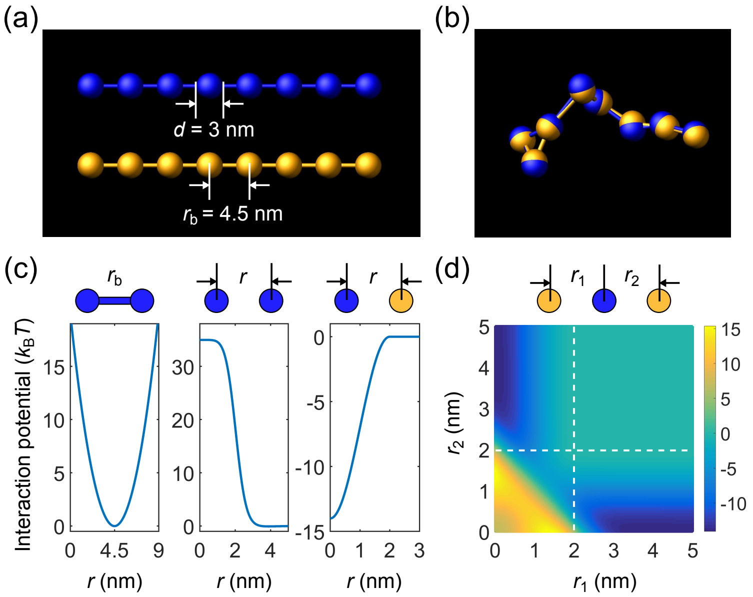


FIG. S1: Coarse-grained molecular-dynamics simulations of two-component multivalent associative polymers. (a) Polymers are modeled as linear chains of spherical particles connected by harmonic bonds. Depicted are $A_8$ (blue) and $B_8$ (yellow). (b) Snapshot of a dimer of $A_8$ and $B_8$ formed in the strong-binding regime. (c) Neighboring monomers in a polymer are connected through a harmonic potential (*left*). Monomers of the same type interact pairwisely through a repulsive potential (*middle*). Monomers of different types interact pairwisely through an attractive potential (*right*). (d) Interaction energy between three particles (one A and two B monomers) as a function of their separation distances. Simultaneous binding of two B monomers to one A monomer is energetically highly disfavored (lower left region) compared to one-to-one binding (dark blue regions).

where $\vec{r}_i$ is the coordinate of particle $i$, $m$ is its mass, $\gamma$ is the friction coefficient, $\vec{f}$ is random thermal noise, and the energy $U(\vec{r}_1, ..., \vec{r}_N)$ contains all interactions between particles, including harmonic bonds, non-specific, and specific interactions (Eqs. S1-S3).

To promote phase equilibrium and ensure that only a single dense condensate is formed, we first initialize the simulation by confining polymers in the region $-50\text{nm} < x < 50\text{nm}$. The attractive interaction between A and B monomers (Eq. S3) is gradually switched on from $U_0 = 0$ to 14 over $10^8$ time steps. The Langevin thermostat is applied using a damping factor $\tau = m/\gamma = 125\text{ns}$, step size $dt = 2.5\text{ns}$, and mass of particle $m = 3534.3\text{ag}$ during this time period. These parameters give the particle the right diffusion coefficient $D = k_B T/(3\pi\eta d)$ for times longer than $\tau$, where $\eta$ is the water viscosity 0.001kg/m/s and $d$ the diameter of the particle. This annealing procedure leads to the formation of a dense phase close to its equilibrated concentration. The confinement is then removed, and the system is equilibrated for $10^8$ more time steps to allow the formation of dilute phase and relaxation of the dense phase. After these procedures, we switch to smaller $\tau = 10\text{ns}$, $dt = 0.5\text{ns}$, and $m = 282.7\text{ag}$ for data recording ($D$ remains the same). The system is relaxed for $10^8$ steps, then the positions of all particles are recorded every $10^6$ steps for 400 recordings. For each choice of valence and stoichiometry, we perform 10 simulation replicates with different random seeds.

To test the effect of finite size on the phase boundaries (Section I D), simulations in Fig. 4 are performed with $N = 5000$ and a box of size 315nm×63nm×63nm, i.e. both total number of monomers and box volume are doubled while the total monomer concentration remains the same (6.64mM). Procedures for equilibration and data recording are the same (including the initial confinement region) except systems are relaxed for $5 \times 10^8$ steps at $dt = 0.5\text{ns}$ before recording, as the larger system requires a longer relaxation time.

#### C. Determining the phase boundaries

To determine the phase boundaries, we need to obtain the concentrations in the dilute and dense phases. The data we recorded for each system contains 4000 snapshots of polymer configurations (10 replicates and 400 time points each). To measure concentrations, we first group polymers into clusters in each snapshot. Connected monomers are grouped into one cluster: two monomers of the same type are connected if they are neighbors in the same polymer, and two monomers of different types are connected if they are within the attraction distance $r_0$=2nm. In most of

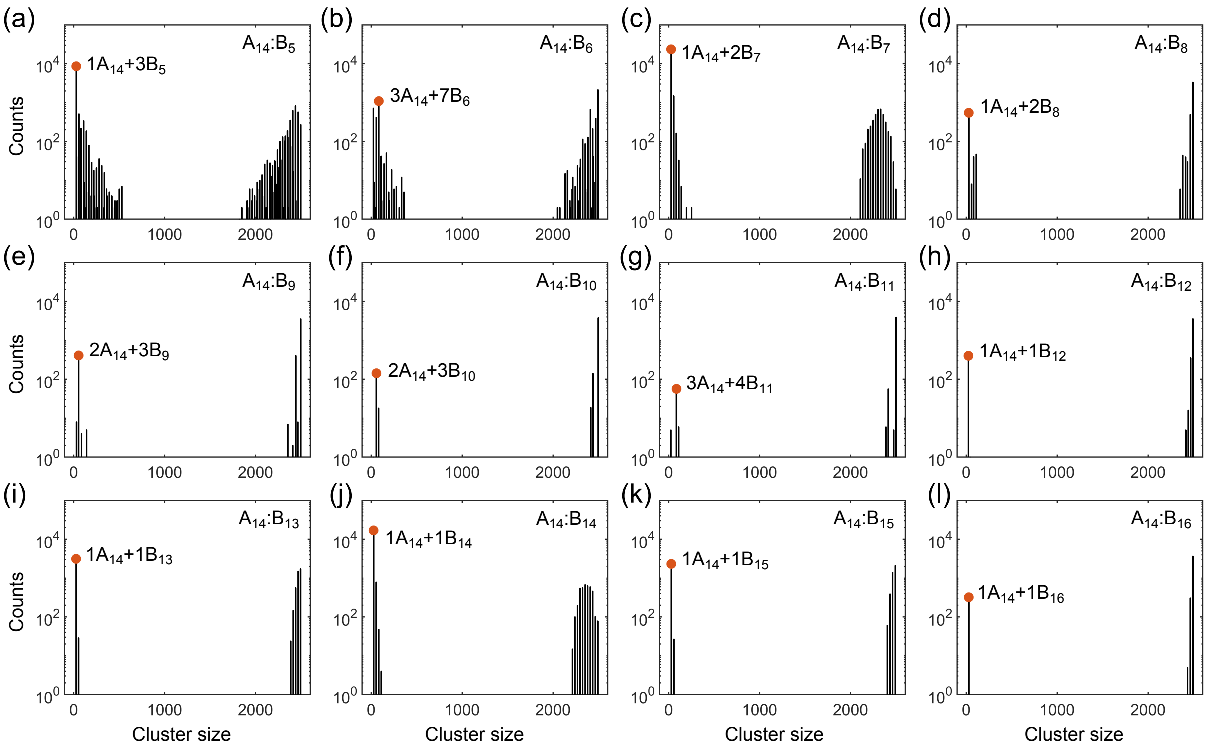


FIG. S2: Histograms of cluster size in monomers in (a-l) $A_{14}$:$B_{5-16}$ systems at equal global monomer stoichiometry. Specific binding strength is $U_0 = 14k_{\mathrm{B}}T$ and total monomer concentration is 6.64mM. Red dots indicate the dominant oligomer in the dilute phase.

TABLE S1: List of all dilute-phase components and monomer percentages for $A_{14}$:$B_{14}$ system at global monomer stoichiometry 1.21.

| Component | $1A_{14}$ | $1A_{14}+1B_{14}$ | $2A_{14}+1B_{14}$ | $2A_{14}+2B_{14}$ | $3A_{14}+2B_{14}$ | $4A_{14}+3B_{14}$ | $5A_{14}+3B_{14}$ |
|---|---|---|---|---|---|---|---|
| Fraction | 0.10% | 37.0% | 7.67% | 0.38% | 19.1% | 16.6% | 0.38% |
| Component | $5A_{14}+4B_{14}$ | $6A_{14}+4B_{14}$ | $6A_{14}+5B_{14}$ | $7A_{14}+5B_{14}$ | $8A_{14}+6B_{14}$ | $10A_{14}+7B_{14}$ | $17A_{14}+13B_{14}$ |
| Fraction | 10.8% | 0.85% | 0.42% | 0.80% | 3.45% | 0.48% | 1.99% |

our simulations, in each snapshot we observe one large cluster which contains most of polymers, and a few to tens of very small clusters (Fig. S2). There is a clear gap between the sizes of these large and small clusters. We define the large cluster as the dense phase, and the smaller clusters as constituents of the dilute phase. In cases where the separation between the dense and dilute phases is unclear, we discard the data set. Figure S2 shows the size distribution of clusters pooled over all snapshots. Table S1 lists all the dilute-phase components and their total monomer percentages in the $A_{14}$:$B_{14}$ system at global monomer stoichiometry 1.21.

To find the dilute- and dense-phase concentrations, we calculate the center of mass of the dense cluster for each snapshot, and recenter the simulation box to this center of mass. We then compute the monomer concentration histogram along the $x$ axis with a bin size 1/50 of box length. The resulting concentration profile has high values in the middle corresponding to the dense-phase concentration, and low values on the two sides corresponding to the dilute-phase concentration. The dilute- and dense-phase concentrations are calculated by averaging the concentration profile over the regions ($x \leq -100$nm or $x \geq 100$nm) and ($-10$nm $\leq x \leq 10$nm), respectively.

#### D. Testing the effect of finite simulation size on the phase boundaries

To test the effect of finite size on the phase boundaries, we compare the dilute- and dense-phase concentrations of systems with $N = 2500$ to systems with $N = 5000$ (Table S2). The systems being compared have the same valence, stoichiometry, and total monomer concentration. Simulations with different total numbers of particles show consistent

TABLE S2: Comparison of dilute- and dense-phase concentrations for systems with different total numbers of particles.

| Valence (Stoichiometry) | $A_{14}$:$B_{14}$ (14:14) | $A_{14}$:$B_{13}$ (14:14) | $A_{14}$:$B_{13}$ (14:13) |
|---|---|---|---|
| $c_{\rm dil}$ (mM) for $N = 2500$ | 0.50±0.06 | 0.08±0.02 | 0.39±0.02 |
| $c_{\rm dil}$ (mM) for $N = 5000$ | 0.45±0.03 | 0.12±0.02 | 0.34±0.03 |
| $c_{\rm den}$ (mM) for $N = 2500$ | 28.0±0.2 | 27.76±0.09 | 25.80±0.08 |
| $c_{\rm den}$ (mM) for $N = 5000$ | 27.93±0.05 | 27.82±0.06 | 25.82±0.06 |

results, suggesting the effect of finite size is minor.

#### E. Measuring the dissociation constants

The phase boundaries of associative polymers from theory are very sensitive to model parameters. Here we extract the dimer and monomer dissociation constants $K_{\rm d}$ and $K_{\rm b}$ from simulations. These values are then utilized in the dimer-gel theory to obtain the free-energy density landscape and predict the phase boundaries. Our simulations are performed in the strong binding regime. At our chosen binding strength of $U_0 = 14k_{\rm B}T$, the long lifetime of each bond means that dimers of valence $\geq 4$ never dissociate in our simulations. This prevents us from directly extracting the dimer dissociation constants for long polymers. We therefore use a reweighting method [3] to obtain the dissociation constant for these dimers.

Briefly, we perform simulations with 1 polymer of type A and 1 polymer of type B of valences $L_1$ and $L_2$ in a cubic box of side $20L_1$nm with periodic boundary conditions. The systems are equilibrated using a Langevin thermostat in the $NVT$ ensemble, all the parameters are the same as the ones used for recording the data in Section I B, except here we use $U_0 = 7k_{\rm B}T$ which is half of the original value, in order to allow dimers to dissociate.

Theoretically, the dissociation constant of a dimer is defined as

$$K_{\rm d}^{-1} = V\frac{\iint_{U_{\rm AB}<0} e^{-\beta U_{\rm A}(\vec{r}_1^{\,\rm A},\ldots,\vec{r}_{L_1}^{\,\rm A})}e^{-\beta U_{\rm B}(\vec{r}_1^{\,\rm B},\ldots,\vec{r}_{L_2}^{\,\rm B})}e^{-\beta U_{\rm AB}(\vec{r}_1^{\,\rm A},\ldots,\vec{r}_{L_1}^{\,\rm A},\vec{r}_1^{\,\rm B},\ldots,\vec{r}_{L_2}^{\,\rm B})}d\vec{r}_1^{\,\rm A}\ldots d\vec{r}_{L_1}^{\,\rm A}d\vec{r}_1^{\,\rm B}\ldots,d\vec{r}_{L_2}^{\,\rm B}}{\iint e^{-\beta U_{\rm A}(\vec{r}_1^{\,\rm A},\ldots,\vec{r}_{L_1}^{\,\rm A})}e^{-\beta U_{\rm B}(\vec{r}_1^{\,\rm B},\ldots,\vec{r}_{L_2}^{\,\rm B})}d\vec{r}_1^{\,\rm A}\ldots d\vec{r}_{L_1}^{\,\rm A}d\vec{r}_1^{\,\rm B}\ldots,d\vec{r}_{L_2}^{\,\rm B}}, \tag{S7}$$

where $\beta = 1/k_{\rm B}T$, and $(\vec{r}_1^{\,\rm A},\ldots,\vec{r}_{L_1}^{\,\rm A})$ and $(\vec{r}_1^{\,\rm B},\ldots,\vec{r}_{L_2}^{\,\rm B})$ are the coordinates of monomers in polymers A and B. $U_{\rm A}(U_{\rm B})$ contain all interactions within the polymer A(B), including bond potentials $U_{\rm b}$ (Eq. S1) and non-specific interactions $U_{\rm r}$ (Eq. S2). $U_{\rm AB}$ contains all the specific interactions between polymers A and B, i.e. the sum of $U_{\rm a}$s (Eq. S3). Integration is over the entire volume $V$. In the denominator, the integration is further confined to the region where $U_{\rm AB} < 0$. Note that for a dissociated dimer $U_{\rm AB} = 0$.

To link the simulation with the definition of $K_{\rm d}$, we define three variables in the simulation:

$$\begin{aligned} &w_1 = e^{\beta E_{\rm AB}}, w_2 = 1, w_3 = e^{-\beta E_{\rm AB}}, \qquad &&\text{if } E_{\rm AB} < 0; \\ &w_1 = w_2 = w_3 = 0, &&\text{if } E_{\rm AB} = 0, \end{aligned} \tag{S8}$$

where $E_{\rm AB} = U_{\rm AB}(\vec{r}_1^{\,\rm A},\ldots,\vec{r}_{L_1}^{\,\rm A},\vec{r}_1^{\,\rm B},\ldots,\vec{r}_{L_2}^{\,\rm B})$. We then have,

$$K_{\rm d}^{-1}(U_0 = 0k_{\rm B}T) = C\iint w_1 e^{-\beta U_{\rm A}}e^{-\beta U_{\rm B}}e^{-\beta U_{\rm AB}}d\vec{r}_1^{\,\rm A}\ldots d\vec{r}_{L_1}^{\,\rm A}d\vec{r}_1^{\,\rm B}\ldots,d\vec{r}_{L_2}^{\,\rm B} = \tilde{C}\langle w_1\rangle, \tag{S9}$$

$$K_{\rm d}^{-1}(U_0 = 7k_{\rm B}T) = C\iint w_2 e^{-\beta U_{\rm A}}e^{-\beta U_{\rm B}}e^{-\beta U_{\rm AB}}d\vec{r}_1^{\,\rm A}\ldots d\vec{r}_{L_1}^{\,\rm A}d\vec{r}_1^{\,\rm B}\ldots,d\vec{r}_{L_2}^{\,\rm B} = \tilde{C}\langle w_2\rangle, \tag{S10}$$

$$K_{\rm d}^{-1}(U_0 = 14k_{\rm B}T) = C\iint w_3 e^{-\beta U_{\rm A}}e^{-\beta U_{\rm B}}e^{-\beta U_{\rm AB}}d\vec{r}_1^{\,\rm A}\ldots d\vec{r}_{L_1}^{\,\rm A}d\vec{r}_1^{\,\rm B}\ldots,d\vec{r}_{L_2}^{\,\rm B} = \tilde{C}\langle w_3\rangle, \tag{S11}$$

where $\langle w_1\rangle$, $\langle w_2\rangle$, and $\langle w_3\rangle$ are the mean values of $w_1$, $w_2$, and $w_3$ obtained by averaging over a simulation with $U_0 = 7k_{\rm B}T$. $C$ and $\tilde{C}$ are constants and are the same in all three equations. Therefore,

$$K_{\rm d}(U_0 = 0k_{\rm B}T)\langle w_1\rangle = K_{\rm d}(U_0 = 7k_{\rm B}T)\langle w_2\rangle = K_{\rm d}(U_0 = 14k_{\rm B}T)\langle w_3\rangle. \tag{S12}$$

On the other hand, it can be shown that the binding probability $\langle w_2\rangle$ for $U_0 = 7k_{\rm B}T$ is

$$\langle w_2\rangle = \frac{\iint w_2 e^{-\beta U_{\rm A}}e^{-\beta U_{\rm B}}e^{-\beta U_{\rm AB}}d\vec{r}_1^{\,\rm A}\ldots d\vec{r}_{L_1}^{\,\rm A}d\vec{r}_1^{\,\rm B}\ldots,d\vec{r}_{L_2}^{\,\rm B}}{\iint e^{-\beta U_{\rm A}}e^{-\beta U_{\rm B}}e^{-\beta U_{\rm AB}}d\vec{r}_1^{\,\rm A}\ldots d\vec{r}_{L_1}^{\,\rm A}d\vec{r}_1^{\,\rm B}\ldots,d\vec{r}_{L_2}^{\,\rm B}} = \frac{1}{1 + K_{\rm d}(U_0 = 7)\left(V - K_{\rm d}^{-1}(U_0 = 0)\right)}. \tag{S13}$$

TABLE S3: List of dissociation constants from simulations

| | $A_1$:$B_1$$^{(t)}$ | $A_1$:$B_1$$^{(d)}$ | $A_1$:$B_1$$^{(r)}$ | $A_2$:$B_2$$^{(d)}$ | $A_2$:$B_2$$^{(r)}$ |
|---|---|---|---|---|---|
| $K_d(U_0 = 0k_BT)$ (mM) | 49.6 | * | 48.9 | * | 12.4 |
| $K_d(U_0 = 7k_BT)$ (mM) | 1.88 | * | 1.90 | * | 0.31 |
| $K_d(U_0 = 14k_BT)$ (mM) | 5.7e-3 | 5.9e-3 | 5.8e-3 | 7.7e-6 | 7.1e-6 |
| | $A_4$:$B_3$$^{(r)}$ | $A_4$:$B_4$$^{(r)}$ | $A_6$:$B_5$$^{(r)}$ | $A_6$:$B_6$$^{(r)}$ | |
| $K_d(U_0 = 0k_BT)$ (mM) | 4.32 | 3.29 | 1.81 | 1.52 | |
| $K_d(U_0 = 7k_BT)$ (mM) | 4.5e-2 | 2.2e-2 | 4.3e-3 | 2.2e-3 | |
| $K_d(U_0 = 14k_BT)$ (mM) | 3.77e-9 | 1.38e-11 | 6.95e-15 | 3.00e-17 | |
| | $A_8$:$B_6$$^{(r)}$ | $A_8$:$B_7$$^{(r)}$ | $A_8$:$B_8$$^{(r)}$ | $A_8$:$B_9$$^{(r)}$ | $A_8$:$B_{10}$$^{(r)}$ |
| $K_d(U_0 = 0k_BT)$ (mM) | 1.15 | 1.01 | 0.90 | 0.79 | 0.73 |
| $K_d(U_0 = 7k_BT)$ (mM) | 9.0e-4 | 4.5e-4 | 2.5e-4 | 1.5e-4 | 1.0e-4 |
| $K_d(U_0 = 14k_BT)$ (mM) | 4.22e-18 | 1.26e-20 | 7.07e-23 | 1.77e-23 | 7.59e-24 |
| | $A_{14}$:$B_{12}$$^{(r)}$ | $A_{14}$:$B_{13}$$^{(r)}$ | $A_{14}$:$B_{14}$$^{(r)}$ | $A_{14}$:$B_{15}$$^{(r)}$ | $A_{14}$:$B_{16}$$^{(r)}$ |
| $K_d(U_0 = 0k_BT)$ (mM) | 0.37 | 0.32 | 0.30 | 0.32 | 0.27 |
| $K_d(U_0 = 7k_BT)$ (mM) | 1.4e-6 | 6.7e-7 | 3.8e-7 | 2.6e-7 | 1.5e-7 |
| $K_d(U_0 = 14k_BT)$ (mM) | 2.74e-35 | 1.08e-37 | 8.85e-40 | 1.80e-40 | 5.16e-41 |

$^{(t)}$ theoretical value, $^{(d)}$ direct simulation, and $^{(r)}$ reweighting method.

Combining Eqs. S12 and S13, we have

$$K_d(U_0 = 0k_BT) = \frac{1 + \langle w_1\rangle - \langle w_2\rangle}{\langle w_1\rangle V}, \tag{S14}$$

$$K_d(U_0 = 7k_BT) = \frac{1 + \langle w_1\rangle - \langle w_2\rangle}{\langle w_2\rangle V}, \tag{S15}$$

$$K_d(U_0 = 14k_BT) = \frac{1 + \langle w_1\rangle - \langle w_2\rangle}{\langle w_3\rangle V}. \tag{S16}$$

Table S3 shows a list of dissociation constants from simulations. The reweighting method provides very accurate monomer-monomer and dimer-dimer dissociation constants, as confirmed by comparing with theory and direct simulations for monomers and polymers of length $L = 2$. The dissociation constants obtained by the reweighting method are used in the dimer-gel theory to predict the phase boundaries (Section II).

### II. THEORETICAL AND NUMERICAL CALCULATIONS

#### A. Derivation of free-energy density for non-specific interactions

The free-energy density due to non-specific interactions can be written as a power expansion in the concentrations [4,5],

$$\frac{F_{ns}}{k_BT} = \frac{1}{2}\sum_{ij} v_{ij}c_ic_j + \frac{1}{6}\sum_{ijk} w_{ijk}c_ic_jc_k, \tag{S17}$$

where the sum is over all the species in the system, including free polymers/monomers, dimers and independent bonds, and $v_{ij}$ and $w_{ijk}$ are two- and three-body interaction parameters.

In the strong-binding regime where the magic-ratio effect is observed, bound monomers strongly overlap, so the size of a bound pair is almost that of a free monomer. For simplicity, we therefore assume the interactions between dimers to be the same as between free polymers of the same type (denoted as $v_d$ and $w_d$), and the interactions between independent bonds to be the same as between free monomers of the same type (denoted as $v_b$ and $w_b$). When independent bonds are preferred (i.e., when $\rho_d \approx 0$), the free-energy density for non-specific interactions is

$$\begin{aligned}\frac{F_{ns}^{ind}}{k_BT} = \; & \frac{v_b}{2}\left[(c_1-c_b)^2+(c_2-c_b)^2+2(c_1-c_b)c_b+2(c_2-c_b)c_b+c_b^2\right]+ \\ & \frac{w_b}{6}\left[(c_1-c_b)^3+(c_2-c_b)^3+3(c_1-c_b)^2c_b+3(c_1-c_b)c_b^2+3(c_2-c_b)^2c_b+3(c_2-c_b)c_b^2+c_b^3\right],\end{aligned} \tag{S18}$$

where $c_1$, $c_2$, and $c_{\rm b}$ are the total concentrations of polymer A, B, and independent bonds in monomeric units. $c_1 - c_{\rm b}$ and $c_2 - c_{\rm b}$ are therefore the concentrations of free monomer A and B. Note that in our simulations, there is no non-specific interaction between free polymers of different types. Therefore, all $v$ and $w$ terms involving different free species are 0. Eq. (S18) simplifies to

$$\frac{F_{\rm ns}^{\rm ind}}{k_{\rm B}T} = \frac{v_{\rm b}}{2}\left(c_1^2 + c_2^2 - c_{\rm b}^2\right) + \frac{w_{\rm b}}{6}\left(c_1^3 + c_2^3 - c_{\rm b}^3\right). \tag{S19}$$

Similarly, when dimers are preferred (i.e., when $c_{\rm b} \approx 0$), the free-energy density for non-specific interactions is

$$\frac{F_{\rm ns}^{\rm dim}}{k_{\rm B}T} = \frac{v_{\rm d}}{2}\left(\rho_1^2 + \rho_2^2 - \rho_{\rm d}^2\right) + \frac{w_{\rm d}}{6}\left(\rho_1^3 + \rho_2^3 - \rho_{\rm d}^3\right), \tag{S20}$$

where $\rho_1$, $\rho_2$, and $\rho_{\rm d}$ are the total concentrations of polymer A, B, and dimers in polymeric units.

As non-specific interactions are only important at high concentrations, we simply set $F_{\rm ns} = F_{\rm ns}^{\rm ind}$. Further, in the strong-binding regime, $c_{\rm b} \approx \min(c_1, c_2)$, so

$$\frac{F_{\rm ns}}{k_{\rm B}T} = \frac{v_{\rm b}}{2}\max(c_1, c_2)^2 + \frac{w_{\rm b}}{6}\max(c_1, c_2)^3. \tag{S21}$$

#### B. Determining model parameters

The developed dimer-gel theory has only four parameters for a given system $A_{L_1}$:$B_{L_2}$: the dissociation constants of dimers and independent bonds $K_{\rm d}$ and $K_{\rm b}$, and the non-specific interaction parameters $v_{\rm b}$ and $w_{\rm b}$. The four parameters together determine the competitiveness of the dilute dimer-phase with the dense gel-phase.

We have extracted the values of $K_{\rm d}$ from simulations (Table. S3). Physically, we expect $K_{\rm b}$ to be close to the monomer-monomer dissociation constant 5.7$\mu$M (Table. S3). It is not exactly the same because the bonds in the condensate are tethered by the backbones of the polymers. We expect the non-specific interaction parameters to be approximately $v_{\rm b} = 2B_2$ and $w_{\rm b} = 3B_3$, where $B_2$ and $B_3$ are the second and third virial coefficients [6]: $v_{\rm b} = 6.8 \times 10^{-2}$mM$^{-1}$ and $w_{\rm b} = 2.2 \times 10^{-3}$mM$^{-2}$ for hard spheres of diameter 3nm (the size of a monomer in the simulations), respectively.

The predicted phase boundaries are sensitive to these parameters. We thus tune $K_{\rm b}$, $v_{\rm b}$, and $w_{\rm b}$ around their estimated values to match the dilute- and dense-phase concentrations of the $A_8$:$B_8$ system from simulation. Specifically, we yield $K_{\rm b} = 3.8 \times 10^{-3}$mM, $v_{\rm b} = 9 \times 10^{-2}$mM$^{-1}$, and $w_{\rm b} = 7 \times 10^{-3}$mM$^{-2}$. These parameters are used for $A_8$:$B_{6\text{-}10}$ systems. Further discussions on how model parameters affect phase boundaries can be found in Section II H.

#### C. Derivation of the transition concentration $c_s$

In the strong-binding regime, the free-energy density contributions from specific interactions in the dimer-dominated and independent bond-dominated limit are, respectively,

$$\frac{F_{\rm s}^{\rm dim}}{k_{\rm B}T} = \rho_- \ln K_{\rm d} + (\rho_+ - \rho_-)\ln\frac{\rho_+ - \rho_-}{e} - \rho_+ \ln\frac{\rho_+}{e}, \tag{S22}$$

$$\frac{F_{\rm s}^{\rm ind}}{k_{\rm B}T} = c_- \ln K_{\rm b} + (c_+ - c_-)\ln\frac{c_+ - c_-}{e} - c_+ \ln\frac{c_+}{e}, \tag{S23}$$

where $\rho_+ = \max(c_1/L_1, c_2/L_2)$, $\rho_- = \min(c_1/L_1, c_2/L_2)$, $c_+ = \max(c_1, c_2)$, and $c_- = \min(c_1, c_2)$. Eqs. (S22) and (S23) are the same as Eqs. (16) and (17).

At given ($c_1$,$c_2$), whether the system will form dimers or independent bonds depends on their relative free energies. For equal valence polymers $L_1 = L_2 = L$, letting $c_+ = sc_-$, Eqs. (S22) and (S23) become

$$\frac{F_{\rm s}^{\rm dim}}{k_{\rm B}T} = \rho_- \ln\frac{(s-1)^{s-1}eK_{\rm d}}{s^s \rho_-}, \tag{S24}$$

$$\frac{F_{\rm s}^{\rm ind}}{k_{\rm B}T} = c_- \ln\frac{(s-1)^{s-1}eK_{\rm b}}{s^s c_-}. \tag{S25}$$

Comparing the two expressions, dimers are favored at low concentrations ($F_{\rm s}^{\rm dim} < F_{\rm s}^{\rm ind}$), and independent bonds are favored at high concentrations ($F_{\rm s}^{\rm ind} < F_{\rm s}^{\rm dim}$). The transition occurs when $F_{\rm s}^{\rm dim} = F_{\rm s}^{\rm ind}$, i.e. at the concentrations

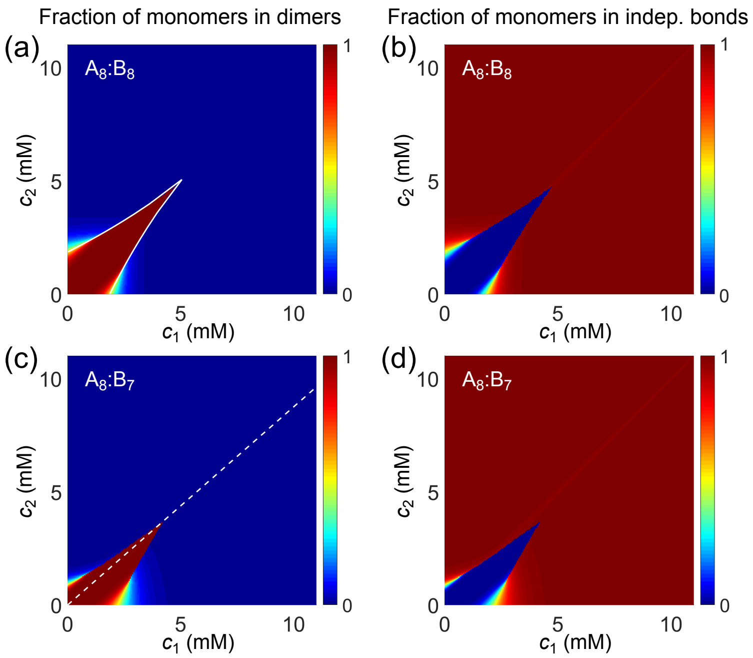


FIG. S3: Fraction of monomers in dimers (a) and in independent bonds (b) for $A_8$:$B_8$ system. White curve in (a) is the transition boundary between dimer- and independent bonds-dominated regions predicted by $c_+(s)$ and $c_-(s)$. Fraction of monomers in dimers (c) and in independent bonds (d) for $A_8$:$B_7$ system. Dashed white line denotes equal polymer stoichiometry.

$c_-(s) = c_0(s-1)^{s-1}s^{-s}$ and correspondingly $c_+(s) = c_0 s(s-1)^{s-1}s^{-s}$ where $c_0 = e(K_{\rm b}^L/(K_{\rm d}L))^{1/(L-1)}$. The boundary between dimer- and independent bond-dominated regions is described by $(c_+(s), c_-(s))$ and $(c_-(s), c_+(s))$, respectively, in the lower and upper halves of the $(c_1, c_2)$ plane (Fig. S3(a), white curve).

#### D. Solving reaction Equations (7) and (8)

The high powers in Eq. (7) and the small value of $K_{\rm d}$ make it difficult to find numerical solutions of $c_{\rm d}$ and $c_{\rm b}$ accurately. To overcome this difficulty, we define a variable $\lambda = c_1 - c_{\rm d1} - c_{\rm b}$ when $\rho_1 <= \rho_2$ and $\lambda = c_2 - c_{\rm d2} - c_{\rm b}$ when $\rho_1 > \rho_2$, and rewrite Eqs. (7) and (8) in terms of $\lambda$. Specifically,

$$\begin{aligned} c_{\rm b} &= \lambda\left(c_2L_1 - c_1L_2 + \lambda L_2\right)\left[K_{\rm b}L_1 + \lambda(L_1 - L_2)\right]^{-1}, \\ c_{\rm d1} &= c_1 - \lambda - c_{\rm b}, \\ K_{\rm d}L_2(c_1 - c_{\rm b} - \lambda)&(1+\lambda K_{\rm b}^{-1})^{L_2-1}(\lambda + c_{\rm b})^{L_1-1} = K_{\rm b}c_{\rm b}\lambda^{L_1-1} \end{aligned} \tag{S26}$$

for $\rho_1 <= \rho_2$, and

$$\begin{aligned} c_{\rm b} &= \lambda\left(c_1L_2 - c_2L_1 + \lambda L_1\right)\left[K_{\rm b}L_2 + \lambda(L_2 - L_1)\right]^{-1}, \\ c_{\rm d2} &= c_2 - \lambda - c_{\rm b}, \\ K_{\rm d}L_1(c_2 - c_{\rm b} - \lambda)&(1+\lambda K_{\rm b}^{-1})^{L_1-1}(\lambda + c_{\rm b})^{L_2-1} = K_{\rm b}c_{\rm b}\lambda^{L_2-1} \end{aligned} \tag{S27}$$

for $\rho_1 > \rho_2$. We then solve Eqs. (S26) and (S27) using the MatLab function *vpasolve* with the constraints $0 < \lambda < c_1$ and $0 < \lambda < c_2$, respectively. *vpasolve* provides all solutions within the specified range. When multiple solutions coexist, we take the one with the lowest $F_{\rm s}$. Numerical solutions of $\rho_{\rm d}$ and $c_{\rm b}$ for $A_8$:$B_8$ and $A_8$:$B_7$ systems are shown in Fig. S3, where the fraction of monomers in dimers is defined as $\rho_{\rm d}/\min(\rho_1, \rho_2)$ and fraction of monomers in independent bonds is defined as $c_{\rm b}/\min(c_1, c_2)$.

#### E. Determining phase boundaries and tie lines

We obtain the free-energy landscape by substituting the numerical solutions of $\rho_{\rm d}$ and $c_{\rm b}$ into Eq. (15) (Fig. S4). We locate the phase boundaries by applying convex-hull analysis to this free-energy landscape using the MatLab

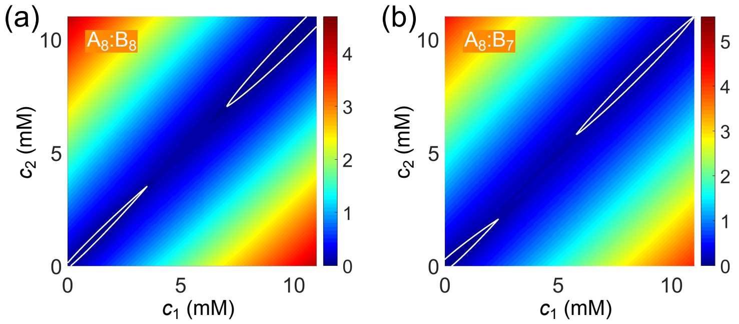


FIG. S4: Free-energy density as a function of global monomer concentrations $(c_1, c_2)$ for (a) $A_8$:$B_8$ and (b) $A_8$:$B_7$ systems. White curves highlight the basins in the dilute dimer-dominated and dense gel-dominated regions.

function *convhull* (Figs. 5(a) and 5(b)).

To find the tie line going through a given initial concentration point $(c_1^{\rm in}, c_2^{\rm in})$, we adopt a modified vector method [7]. We first draw a line though this point along a direction defined by an angle $\alpha$, and then find the crossing points between this line and the phase boundaries, i.e. $(c_1^{\rm dil}, c_2^{\rm dil})$ and $(c_1^{\rm den}, c_2^{\rm den})$. The "true" $\alpha$ is the one that minimizes the free-energy density of mixing $pF(c_1^{\rm dil}, c_2^{\rm dil}) + (1-p)F(c_1^{\rm den}, c_2^{\rm den})$, where $p = r^{\rm den}/(r^{\rm dil} + r^{\rm den})$ is the volume fraction of dilute phase and $1-p$ is the volume fraction of dense phase, and $r^{\rm dil}$ and $r^{\rm den}$ are the distances between the point $(c_1^{\rm in}, c_2^{\rm in})$ and the two points $(c_1^{\rm dil}, c_2^{\rm dil})$ and $(c_1^{\rm den}, c_2^{\rm den})$, respectively.

In order to compare with the simulated dilute- and dense-phase concentrations, we use the initial concentrations from simulations for the specified system at given stoichiometry $s$: $c_1 = c_t s/(1+s)$ and $c_2 = c_t/(1+s)$, where the total monomer concentration is $c_t = 6.64$mM. We then find the tie line going through this initial concentration point and the corresponding $(c_1^{\rm dil}, c_2^{\rm dil})$ and $(c_1^{\rm den}, c_2^{\rm den})$. For simplicity, we show the total dilute- and dense-phase concentrations $c^{\rm dil} = c_1^{\rm dil} + c_2^{\rm dil}$ and $c^{\rm den} = c_1^{\rm den} + c_2^{\rm den}$ in Fig. 5(c) and 5(d).

#### F. A simple approximation for the free-energy density $F$

Finding the numerical solutions of Eqs. (7) and (8) becomes difficult with increasing valence. From the full solutions of $c_{\rm d}$ and $c_{\rm b}$ (Fig. S3), we see that in the dimer-dominated region $c_{\rm b}$ is almost 0, and in the independent bond-dominated region $c_{\rm d}$ is almost 0. Therefore, we can approximate the free-energy density as the lower value of the two limiting cases

$$F = F_{\rm ni} + \min(F_{\rm s}^{\rm dim}, F_{\rm s}^{\rm ind}) + F_{\rm ns}, \tag{S28}$$

where

$$\frac{F_{\rm s}^{\rm dim}}{k_{\rm B}T} = \rho_{\rm d}\ln K_{\rm d} + \rho_{\rm d}\ln\frac{\rho_{\rm d}}{e} + (\rho_1-\rho_{\rm d})\ln\frac{\rho_1-\rho_{\rm d}}{e} + (\rho_2-\rho_{\rm d})\ln\frac{\rho_2-\rho_{\rm d}}{e} - \rho_2\ln\frac{\rho_2}{e} - \rho_1\ln\frac{\rho_1}{e}, \tag{S29}$$

$$\frac{F_{\rm s}^{\rm ind}}{k_{\rm B}T} = c_{\rm b}\ln K_{\rm b} + c_{\rm b}\ln\frac{c_{\rm b}}{e} + (c_1-c_{\rm b})\ln\frac{c_1-c_{\rm b}}{e} + (c_2-c_{\rm b})\ln\frac{c_2-c_{\rm b}}{e} - c_1\ln\frac{c_1}{e} - c_2\ln\frac{c_2}{e}, \tag{S30}$$

and

$$\rho_{\rm d} = \frac{1}{2}\left[\rho_1+\rho_2+K_{\rm d} - \sqrt{(\rho_1+\rho_2+K_{\rm d})^2 - 4\rho_1\rho_2}\right], \tag{S31}$$

$$c_{\rm b} = \frac{1}{2}\left[c_1+c_2+K_{\rm b} - \sqrt{(c_1+c_2+K_{\rm b})^2 - 4c_1c_2}\right] \tag{S32}$$

are solutions of Eqs. (9) and (10). Eq. (S28) provides a very good approximation to the full expression for $F$ (Eq. (15)). The phase diagrams derived from Eqs. (15) and (S28) are almost identical for $A_{14}$:$B_{14}$ (Fig. S5). Results in Fig. S6 are obtained with this approximation (Eqs. (S28-S32)) as there are difficulties solving Eqs. (7) and (8) numerically for $A_{14}$:$B_{12-16}$ systems due to their high valences.

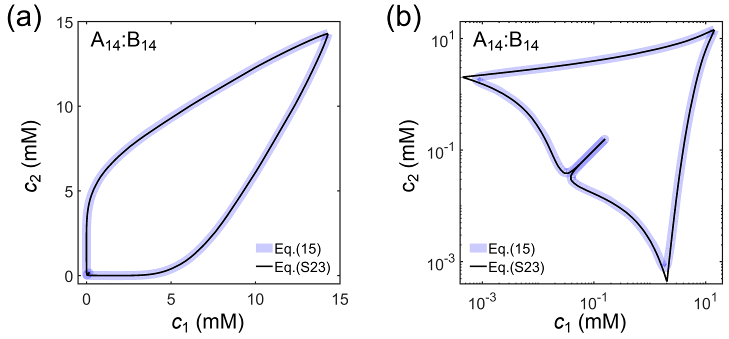


FIG. S5: Comparison of phase diagrams derived from the full expression for $F$ (Eq. (15)) and the approximate expression (Eq. (S28)) shown on a (a) linear and (b) log scale for $A_{14}$:$B_{14}$ system.

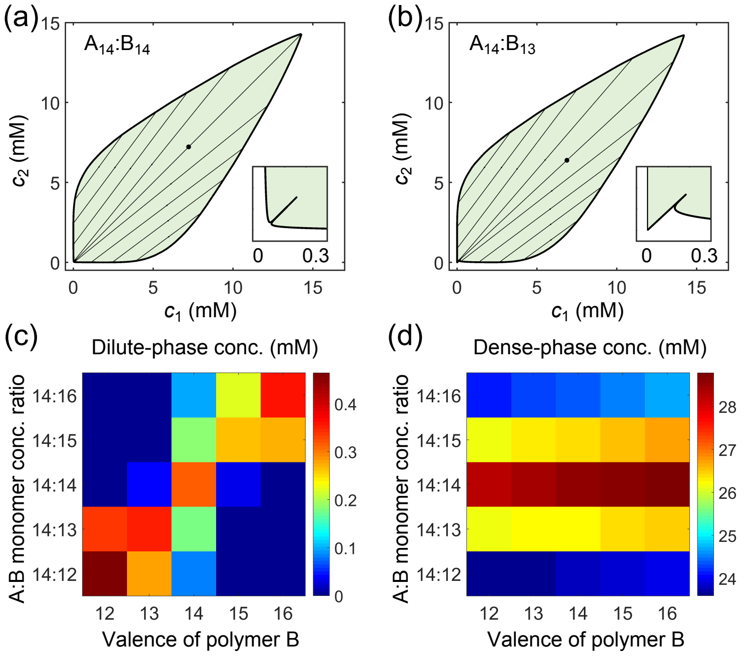


FIG. S6: A dimer-gel theory predicts the generalized magic-number effect. Phase diagrams of (a) $A_{14}$:$B_{14}$ and (b) $A_{14}$:$B_{13}$ systems: one-phase region white, two-phase region green. The dilute- and dense-phase concentrations are connected by representative tie lines. The tie line along the direction of equal polymer stoichiometry is denoted with a black dot. Inset: enlarged dilute-phase boundaries. Monomer concentrations in (c) dilute and (d) dense phases for systems $A_{14}$:$B_{12\text{-}16}$ at global monomer stoichiometries 14:12-16 and total monomer concentration 6.64mM. Parameters: $v_{\mathrm{b}} = 7 \times 10^{-2}\mathrm{mM}^{-1}$, $w_{\mathrm{b}} = 5 \times 10^{-3}\mathrm{mM}^{-2}$, $K_{\mathrm{b}} = 3.8 \times 10^{-3}\mathrm{mM}$, and $K_{\mathrm{d}}$ in Table S3.

#### G. Correlated binding in the dense phase

In the dimer-gel theory, we assume that monomers of different types can associate independently in the dense phase. However, as monomers belonging to the same polymer are tethered together, neighboring monomers in one polymer are more likely to bind to neighboring monomers in another polymer, i.e. there are correlations in binding. To quantify this correlation, we first identify consecutive segments in a polymer that bind to consecutive segments in another polymer. (For example, if in polymer 1 of type A, monomers 1, 2, 3, and 4 bind to monomers 2, 4, 3, and 5 of one polymer of type B, monomer 5 binds to monomer 8 of a second polymer of type B, and monomers 6, 7, and 8 bind to monomers 1, 2, and 3 of a third polymer of type B, then there are three individual segments in polymer 1 of type A with lengths 4, 1, and 3.) Clearly, what should be considered to be "independent" are not individual monomers but rather these consecutively bound segments. To quantify the length of these segments, we measure the probability $p(l)$ that a bound monomer is in a segment of length $l$. Figure S7 shows the probability distribution $p(l)$ for simulated $A_8$:$B_8$ and $A_{14}$:$B_{14}$ systems at equal stoichiometry. The mean length of "independent" segments is 1.8

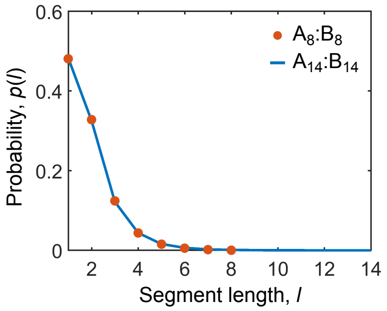


FIG. S7: Binding between monomers of different types in the dense phase is correlated. Probability distribution $p(l)$ for finding a bound monomer to be in a consecutively bound segment of length $l$ in the dense phase of $A_8$:$B_8$ and $A_{14}$:$B_{14}$ systems at equal stoichiometry.

for both cases.

#### H. Effects of model parameters on phase boundaries

The dimer-gel theory has only a handful of parameters: the valences $L_1$ and $L_2$ of polymers A and B, the dissociation constants $K_\mathrm{d}$ and $K_\mathrm{b}$ of dimers and independent bonds, and the non-specific interaction parameters $v_\mathrm{b}$ and $w_\mathrm{b}$. These parameters together determine the competitiveness of the dilute dimer-phase with the dense gel-phase. We first fix $K_\mathrm{b} = 3.8 \times 10^{-3}$mM, $v_\mathrm{b} = 9 \times 10^{-2}$mM$^{-1}$, and $w_\mathrm{b} = 7 \times 10^{-3}$mM$^{-2}$ (see Section II B for how these parameters are derived, and note that the values of $K_\mathrm{d}$ are taken directly from simulations (Table S3)), and explore the dependence of phase boundaries on the valences $L_1$, $L_2$ and on the stoichiometry.

Figures S8(a) and S8(b) show phase diagrams and dilute-phase concentrations for $A_4$:$B_4$ to $A_{14}$:$B_{14}$ systems. For

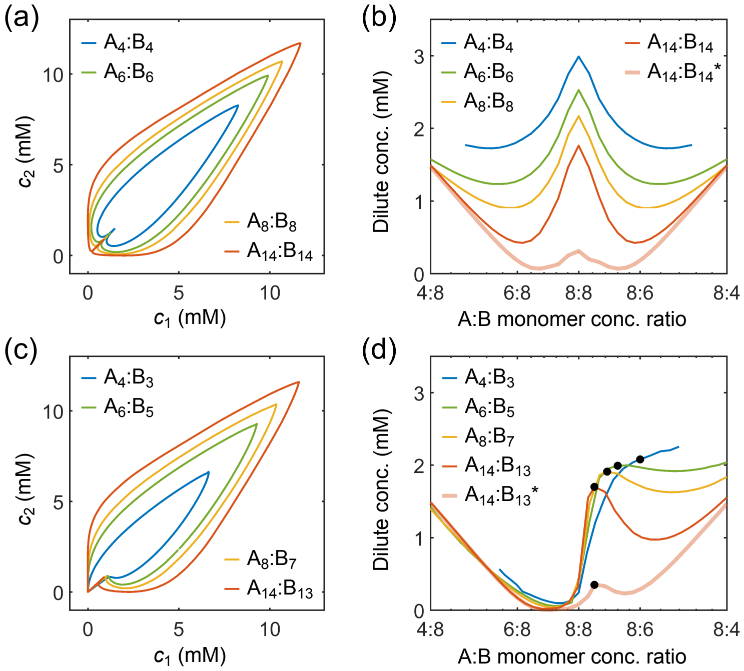


FIG. S8: Effects of model parameters on the phase boundaries in the strong-binding regime. (a) Phase diagrams and (b) dilute-phase concentrations at different global stoichiometries for $A_4$:$B_4$ to $A_{14}$:$B_{14}$ systems. (c) Phase diagrams and (d) dilute-phase concentrations at different global stoichiometries for $A_4$:$B_3$ to $A_{14}$:$B_{13}$ systems. Black dots indicate equal *polymer* stoichiometries. Parameters: $K_\mathrm{b} = 3.8 \times 10^{-3}$mM, values of $K_\mathrm{d}$ in Table S3 for binding strength $U_0 = 14k_\mathrm{B}T$, $v_\mathrm{b} = 9 \times 10^{-2}$mM$^{-1}$ and $w_\mathrm{b} = 7 \times 10^{-3}$mM$^{-2}$ for all systems except for $A_{14}$:$B_{14}^*$ and $A_{14}$:$B_{13}^*$ where $v_\mathrm{b} = 7 \times 10^{-2}$mM$^{-1}$ and $w_\mathrm{b} = 5 \times 10^{-3}$mM$^{-2}$. Total monomer concentration is 6.64 mM.

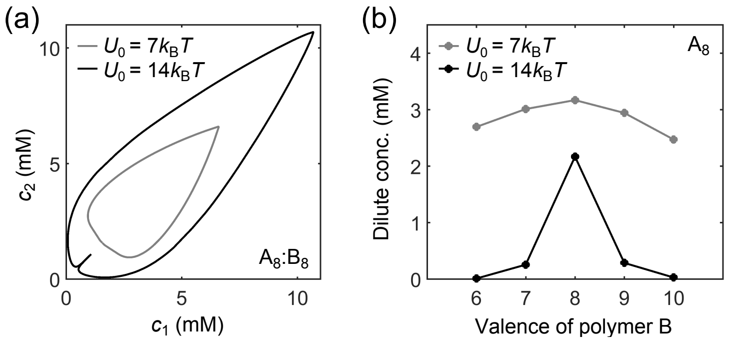


FIG. S9: The magic-ratio effect disappears in the weak-binding regime. (a) Phase diagrams for $A_8$:$B_8$ system with different binding strengths $U_0 = 7k_BT$ (gray) and $U_0 = 14k_BT$ (black). (b) Dilute-phase concentrations for $A_8$:$B_{6-10}$ systems at equal global monomer stoichiometry with different binding strengths $U_0 = 7k_BT$ (gray) and $U_0 = 14k_BT$ (black). Parameters: $K_b = 1.88$mM for $U_0 = 7k_BT$, $K_b = 3.8 \times 10^{-3}$mM for $U_0 = 14k_BT$, values of $K_d$ in Table S3, $v_b = 9 \times 10^{-2}$mM$^{-1}$, and $w_b = 7 \times 10^{-3}$mM$^{-2}$. Total monomer concentration is 6.64 mM.

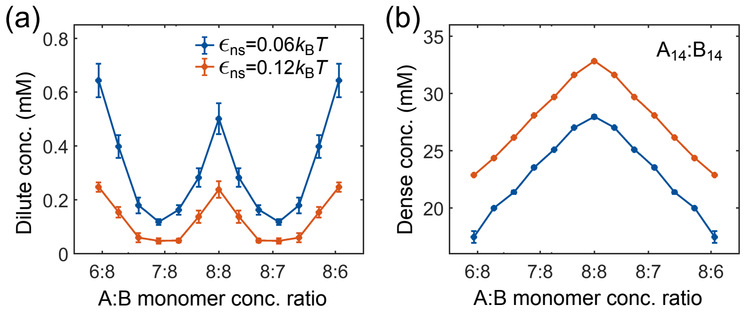


FIG. S10: Strength of non-specific interactions strongly influences phase boundaries in simulations. Monomer concentrations in (a) dilute and (b) dense phases for $A_{14}$:$B_{14}$ system at different global stoichiometries with non-specific attraction tail strength $\epsilon_{ns} = 0.06k_BT$ (blue) and $\epsilon_{ns} = 0.12k_BT$ (red). Specific interaction strength $U_0 = 14k_BT$ and total monomer concentration is 6.64 mM.

these equal valence systems, the dilute-/dense-phase concentrations decreases/increases with increasing valence, and the magic-number effect with respect to stoichiometry is enhanced with increasing valence in terms of dilute-phase peak-to-valley ratio. Figures S8(c) and S8(d) show phase diagrams and dilute-phase concentrations for $A_4$:$B_3$ to $A_{14}$:$B_{13}$ systems. Note that the shape of the dilute phase boundary transitions from a shoulder to a peak with increasing valence. All these features are consistent with the simulation results in Fig. 3. Figure S9(b) shows the dilute-phase concentrations for $A_8$:$B_{6-10}$ systems at equal monomer stoichiometry. The dilute-phase concentration is sharply peaked at $A_8$:$B_8$, consistent with the simulation results in Fig. 2.

However, the dilute-phase concentration of $A_{14}$:$B_{14}$ does not quantitatively agree with simulation results (Fig. S8(b) vs. Fig. 3(a)). Also, the dilute-phase concentrations for unequal valence systems do not decrease with increasing valence (Fig. S8(d) vs. Fig. 3(c)). What is the origin of these discrepancies? Intuitively, the dense phase properties are determined by $K_b$, $v_b$, and, $w_b$. Using the same values of these parameters for systems with different valences means that we are treating the dense phases of these systems as exactly equivalent. However, there are more polymer backbone bonds in higher valence systems. These backbone bonds, from a mean-field point of view, act like attractive potentials between monomers, which effectively reduces the non-specific repulsion between monomers. Therefore, the dense phases of higher valence systems are energetically favored, and we expect correspondingly lower values of $v_b$ and $w_b$ for valence 14 systems compared to valence 8 systems. Indeed, we find that somewhat smaller non-specific interaction parameters, $v_b = 7 \times 10^{-2}$mM$^{-1}$ and $w_b = 5 \times 10^{-3}$mM$^{-2}$, lead to quantitative agreement of the dilute- and dense-phase boundaries for $A_{14}$:$B_{14}$ and $A_{14}$:$B_{13}$ systems with the simulation results (Figs. S8(b) and S8(d), curves labeled with *, compared to Figs. 3(a) and 3(c)). We thus use these parameters for $A_{14}$:$B_{12\text{-}16}$ systems in Fig. S6. Simulations confirm that lower non-specific repulsion leads to decreased dilute-phase concentration and increased dense-phase concentration (Fig. S10).

Finally, the dimer-gel theory also predicts that the magic-ratio effect in the weak-binding regime (Fig. S9(a) and S9(b), gray curves), consistent with the simulation results (Fig. 2(a)).

[1] S. Plimpton, J. Comput. Phys. **117**, 1 (1995).
[2] P. Langevin, C. R. Acad. Sci. **146**, 530 (1908).
[3] D. Frenkel and B. Smit, *Understanding Molecular Simulation: from Algorithms to Applications*, vol. 1 (Elsevier, 2001).
[4] A. N. Semenov and M. Rubinstein, Macromolecules **31**, 1373 (1998).
[5] P.-G. De Gennes, *Scaling Concepts in Polymer Physics* (Cornell university press, 1979).
[6] S. Katsura, Phys. Rev. **115**, 1417 (1959).
[7] A. Marcilla, J. A. Labarta, M. D. Serrano Cayuelas, and M. d. M. Olaya *et al.* (2011).
[8] See LAMMPS manual at https://lammps.sandia.gov/doc/Manual.html for details about this potential.
